# Switch-like Gene Expression Modulates Disease Susceptibility

**DOI:** 10.1101/2024.08.24.609537

**Authors:** Alber Aqil, Yanyan Li, Zhiliang Wang, Saiful Islam, Madison Russell, Theodora Kunovac Kallak, Marie Saitou, Omer Gokcumen, Naoki Masuda

## Abstract

A fundamental challenge in biomedicine is understanding the mechanisms predisposing individuals to disease. While previous research has suggested that switch-like gene expression is crucial in driving biological variation and disease susceptibility, a systematic analysis across multiple tissues is still lacking. By analyzing transcriptomes from 943 individuals across 27 tissues, we identified 1,013 switch-like genes. We found that only 31 (3.1%) of these genes exhibit switch-like behavior across all tissues. These universally switch-like genes appear to be genetically driven, with large exonic genomic structural variants explaining five (∼18%) of them. The remaining switch-like genes exhibit tissue-specific expression patterns. Notably, tissue-specific switch-like genes tend to be switched on or off in unison within individuals, likely under the influence of tissue-specific master regulators, including hormonal signals. Among our most significant findings, we identified hundreds of concordantly switched-off genes in the stomach and vagina that are linked to gastric cancer (41-fold, *p*<10^-4^) and vaginal atrophy (44-fold, *p*<10^-4^), respectively. Experimental analysis of vaginal tissues revealed that low systemic levels of estrogen lead to a significant reduction in both the epithelial thickness and the expression of the switch-like gene *ALOX12*. We propose a model wherein the switching off of driver genes in basal and parabasal epithelium suppresses cell proliferation therein, leading to epithelial thinning and, therefore, vaginal atrophy. Our findings underscore the significant biomedical implications of switch-like gene expression and lay the groundwork for potential diagnostic and therapeutic applications.

## Introduction

The study of gene expression began in earnest with the characterization of lactose-metabolizing switch-like genes in *E. coli* ^*1*^. The presence of lactose triggered the production of enzymes needed to metabolize it, while these enzymes were absent when lactose was not present. These genes acted like switches, toggling between “on” and “off” states based on the presence or absence of lactose, respectively. In subsequent decades, the discovery of enhancer elements ^2–4^, epigenetic modifications ^5–8^, and transcription factor dynamics ^9^ revealed that gene expression in humans is more nuanced, resembling a dimmer more often than a simple on-and-off mechanism. Consequently, the study of switch-like genes in humans was largely relegated to the narrow realm of Mendelian diseases ^10–12^.

The recent availability of population-level RNA-sequencing data from humans has made it possible to systematically identify switch-like versus dimmer-like genes. For dimmer-like genes in a given tissue, we expect expression levels across individuals to be continuously distributed with a single mode, i.e., a unimodal distribution. In contrast, expression levels of switch-like genes in a given tissue are expected to exhibit a bimodal distribution, with one mode representing the “off” state and the other representing the “on” state. As we will detail, bimodal expression across individuals is a characteristic of a gene in a specific tissue, referred to as a gene-tissue pair. We define a gene as switch-like if it exhibits bimodal expression in at least one tissue. Most of the recent studies on bimodal gene expression are related to cancer biology, associating on and off states to different disease phenotypes and their prognoses ^13–15^. These cancer studies have already produced promising results for personalized medicine ^16^. However, to our knowledge, the only study focusing on switch-like genes in non-cancerous tissues across individuals restricted their analysis to muscle tissue ^17^. As a result, the dynamics of switch-like expression across the multi-tissue landscape remain unknown. We hypothesize that switch-like expression is ubiquitous but often tissue-specific. We further hypothesize that these tissue-specific expression trends underlie common disease states. Therefore, the analysis of switch-like genes across tissues and individuals may provide a means for early diagnosis and prediction of human disease.

Here, we systematically identified switch-like genes across individuals in 27 tissues. Our results explain the regulatory bases of switch-like expression in humans, highlighting genomic structural variation as a major factor underlying correlated switch-like expression in multiple tissues. Furthermore, we identified groups of switch-like genes in the stomach and vagina for which the “off” state predisposes individuals to gastric cancer and vaginal atrophy, respectively. Overall, these findings improve our understanding of the regulation of switch-like genes in humans. They also suggest promising future paths for preventative biomedical interventions.

## RESULTS

### Tissue-specificity of bimodal expression

The misregulation of highly expressed genes often has consequences for health and fitness. To systematically identify biomedically relevant switch-like genes in humans, we focused on 19,132 genes that are highly expressed (mean TPM > 10) in at least one of the 27 tissues represented in the GTEx database (**Figure 1A; Figure 1B; Table S1**). For each of the 516,564 gene-tissue pairs (19,132 genes x 27 tissues), we applied the dip test of unimodality ^18^ to the expression level distribution across individuals (**Figure 1C**). Employing the Bejamini-Hochberg procedure for multiple hypotheses correction, we identified 1,013 switch-like genes (**Figure 1C; Methods; Table S2**). The expression of these genes is bimodally distributed in at least one tissue, such that it is switched “off” for one subset of individuals and switched “on” for the rest of the individuals.

**Figure 1.**
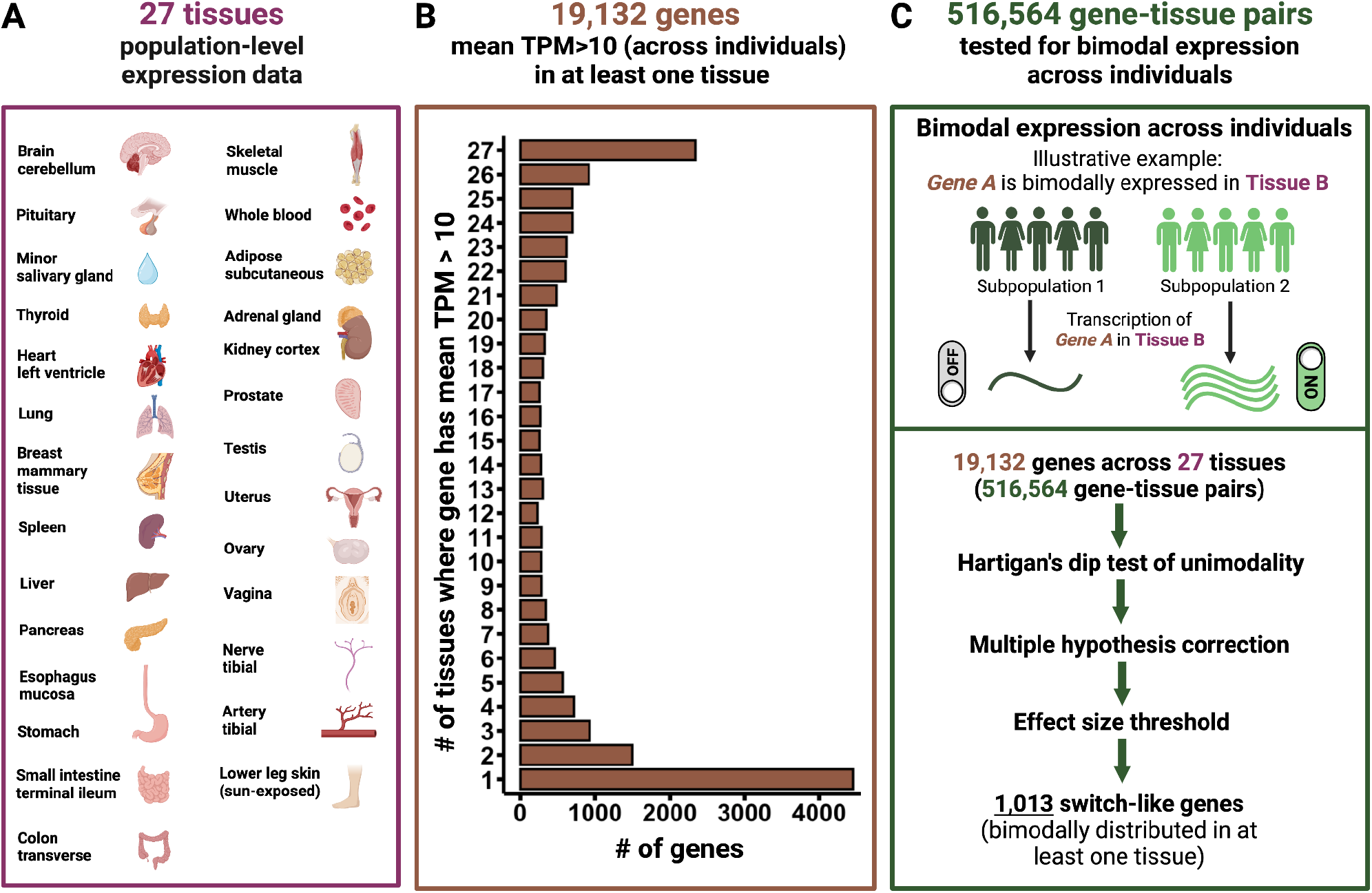
Methodological framework. **A**. List of 27 tissues used in this study. **B**. Distribution of 19,132 genes by the number of tissues in which they are highly expressed. **C**. Bimodal expression is a property of a gene-tissue pair. We tested 516,564 gene-tissue pairs (19,132 genes x 27 tissues) for bimodal expression across individuals. When a gene-tissue pair exhibits switch-like (bimodal) expression, the individuals divide into two subpopulations: one with the gene switched off, and the other with the gene switched on.

Expression of different switch-like genes may be bimodally distributed in different numbers of tissues. We contend that genes that are bimodally expressed across all tissues are likely so due to a germline genetic polymorphism driving switch-like expression across tissues. If this is the case, the expression of these genes would be highly correlated across pairs of tissues. Given this insight, discovering universally bimodal genes is more tractable using tissue-to-tissue co-expression of each gene. Therefore, for each gene, we calculated the pairwise correlation of expression levels across pairs of tissues (**Methods; Table S3**). To visualize tissue-to-tissue co-expression patterns of genes, we performed principal component analysis (PCA) on the tissue-to-tissue gene co-expression data (**Table S4**). We emphasize that we are referring to the co-expression of the same gene across pairs of tissues instead of the co-expression of pairs of genes in the same tissue. In the space spanned by the first two principal components (explaining 35.3% and 3.47% of the variance, respectively), switch-like genes form two major clusters (cluster 1 and cluster 2**; Methods**), dividing along PC1 (**Figure 2A**). Applying PCA exclusively to switch-like genes reveals the further division of cluster 2 into two distinct subclusters – cluster 2A and cluster 2B – in the space spanned by the first two principal components (explaining 58.1% and 4.25% of the variance, respectively) (**Figure 2B; Table S5**).

**Figure 2.**
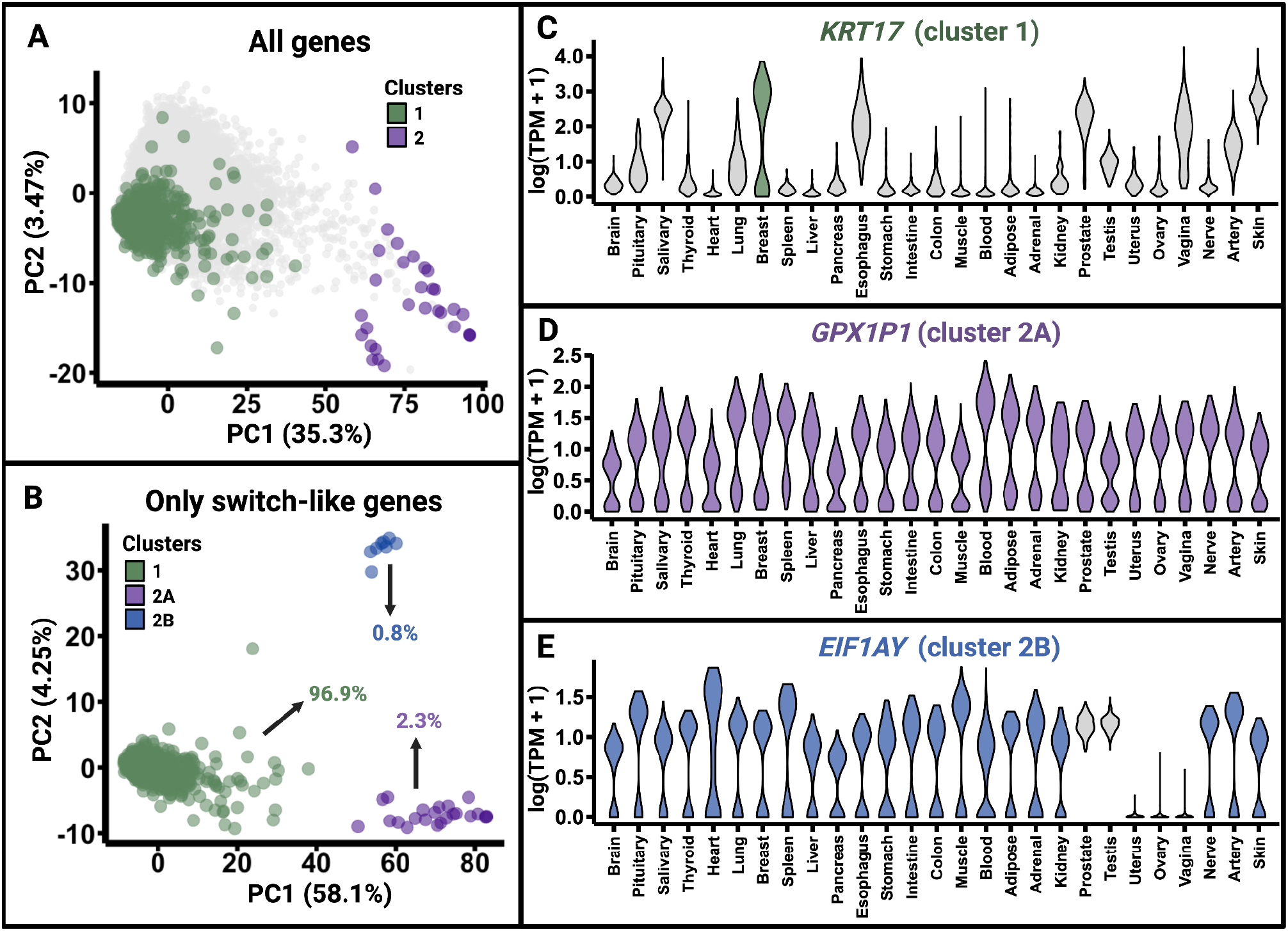
Categorization of switch-like genes. **A**. PCA analysis of tissue-pair correlations of gene expression. Each point represents a gene. When we perform PCA on the tissue-to-tissue co-expression vectors for 19,132 genes, the switch-like genes divide into two clusters. Cluster 1 primarily represents genes that are bimodally expressed in a tissue-specific manner, while cluster 2 represents genes that are bimodally expressed in at least all non-sex-specific tissues. **B**. Performing PCA on the co-expression vectors of only switch-like genes further divides cluster 2 into two subclusters: cluster 2A, which contains genes that are bimodally expressed across all 27 tissues, and cluster 2B, which contains genes that are bimodally expressed in all 22 tissues common to both sexes, but not in the five sex-specific tissues. **C-E**. Violin plots display the expression levels in all 27 tissues for representative genes from cluster 1, cluster 2A, and cluster 2B, respectively.

Manual inspection reveals that cluster 1, which contains 954 genes, represents genes, such as *KRT17*, with bimodal expression in a small subset of tissues (**Figure 2C**). Cluster 2A consists of 23 genes, such as *GPX1P1*, with bimodal expression in all tissues (**Figure 2D**). Lastly, cluster 2B represents eight genes, such as *EIF1AY*, with bimodal expression in all non-sex-specific tissues but not in sex-specific tissues (**Figure 2E**). We will refer to genes in cluster 1 as “tissue-specific switch-like genes.” Although some of them are bimodally expressed in more than one tissue, these genes tend to exhibit high tissue specificity in their bimodal expression. Genes in cluster 2 will be referred to as “universally switch-like genes.”

### Genetic variation underlies universally switch-like genes

We found that 3.1% of all switch-like genes (i.e., the proportion of switch-like genes that are in cluster 2) show clear bimodal expression, at least in all tissues common to both sexes. We contend that germline genetic variation across individuals likely underlies the universally switch-like gene expression, specifically due to four major types of genetic variants. Firstly, we expect genes on the Y chromosome to show bimodal expression in all tissues common to both sexes since these genes are present in males and absent in females (**Figure 3A**). Consistent with this reasoning, seven out of the eight genes in cluster 2B lie within the male-specific region of the Y-chromosome ^19^; the remaining gene in cluster 2B is *XIST*, showing female-specific expression. Secondly, a homozygous gene deletion would result in the gene being switched off (**Figure 3B**). We found five such genes in cluster 2A for which genomic structural variants likely underlie the observed universally switch-like expression; four genes are affected by gene deletions, and the remaining one by an insertion into the gene. Thirdly, the homozygous deletion of a regulatory element can also switch off a gene (**Figure 3C**). While we did not find any examples of this scenario, it remains a theoretical possibility. Lastly, a loss-of-function single nucleotide variant (SNV) or short indel, which disrupts gene function, can switch off the gene (**Figure 3D**). We identified five genes in cluster 2A where such SNVs cause universal bimodality.

**Figure 3.**
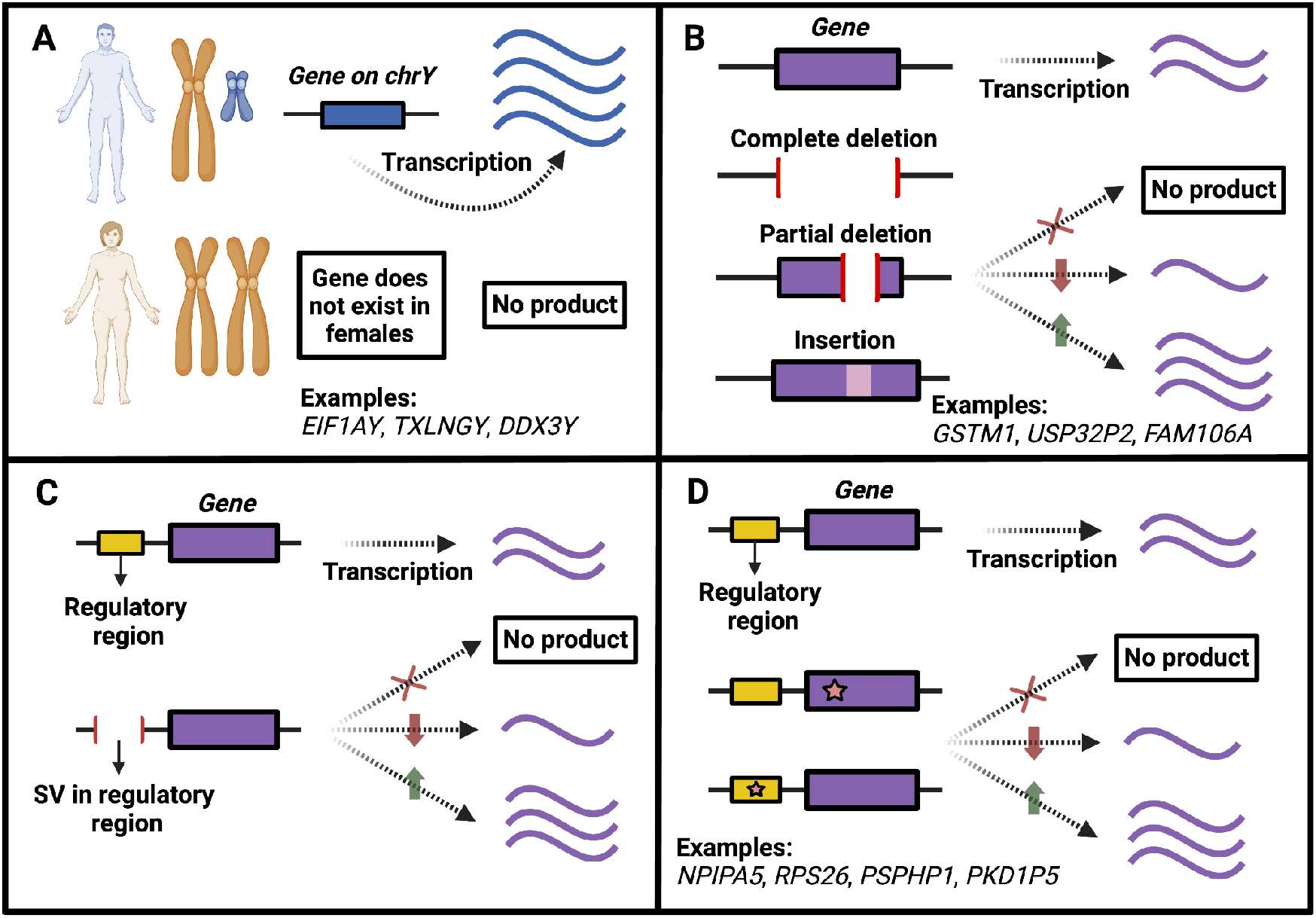
Genetic bases of universally switch-like gene expression (cluster 2). **A**. Genes on the Y chromosome are expressed only in males, leading to bimodal expression in non-sex-specific tissues. **B**. Common structural variants, such as deletions or insertions, may lead to increased, decreased, or no expression in all tissues relative to individuals who carry the alternative allele. **C**. Common structural variants affecting a genomic region regulating a gene may lead to increased, decreased, or no expression in all tissues, relative to individuals who carry the alternative allele. **D**. Common single nucleotide variants or short indels affecting a gene or its regulatory region may lead to increased, decreased, or no expression in all tissues relative to individuals who carry the alternative allele.

Remarkably, we could genetically explain the expression of 10 out of 23 (43%) cases in cluster 2A despite the small number of genes fitting our conservative definition for universally switch-like genes. SNVs underlie five of these cases (**Figure 3B**), while structural variants underlie the remaining five cases (**Figure 3D)**. Thus, out of the 10 cases where we can explain the genetic underpinnings of switch-like expression, 50% involve genomic structural variation, highlighting the importance of this type of genetic variation. Although we could not identify the genetic variation underlying the bimodal expression of the remaining 13 genes in cluster 2A, their consistent and highly correlated switch-like expression across all tissues strongly suggests a genetic basis. We anticipate that better resolution assemblies and detailed regulatory sequence annotations will help identify the genetic variants responsible for the remaining universally switch-like genes.

We highlight a clear example of a common structural variant leading to universally switch-like expression (**Figure 3B**). *USP32P2* and *FAM106A* – both universally switch-like genes – are bimodally expressed in all 27 tissues. Both genes show high levels of tissue-to-tissue co-expression. A common 46 kb deletion (esv3640153), with a global allele frequency of ∼25%, completely deletes both genes (**Figure 4A-B**). We propose that this deletion accounts for the universal switch-like expression of both *USP32P2* and *FAM106A* in all tissues. For illustration, we show the expression level distributions of *USP32P2* and *FAM106A* in the cerebellum (**Figures 4C-D**). Indeed, the haplotype harboring this deletion is strongly associated with the downregulation of both genes in all 27 tissues (*p*<10^-5^ for every single gene-tissue pair, **Methods**). We note that the under-expression of *USP32P2* in sperm is associated with male infertility ^20^, and plausibly, homozygous males for the deletion may be prone to infertility. Additionally, *FAM106A* interacts with SARS-CoV-2 and is downregulated after infection, at least in lung-epithelial cells ^21–23^. Individuals with *FAM106A* already switched off may develop more severe COVID-19 symptoms upon infection, though further investigation is needed. The case of *FAM106A* and *USP32P2* exemplifies the link between disease and bimodal gene expression, a theme we will explore further in the remainder of this text.

**Figure 4.**
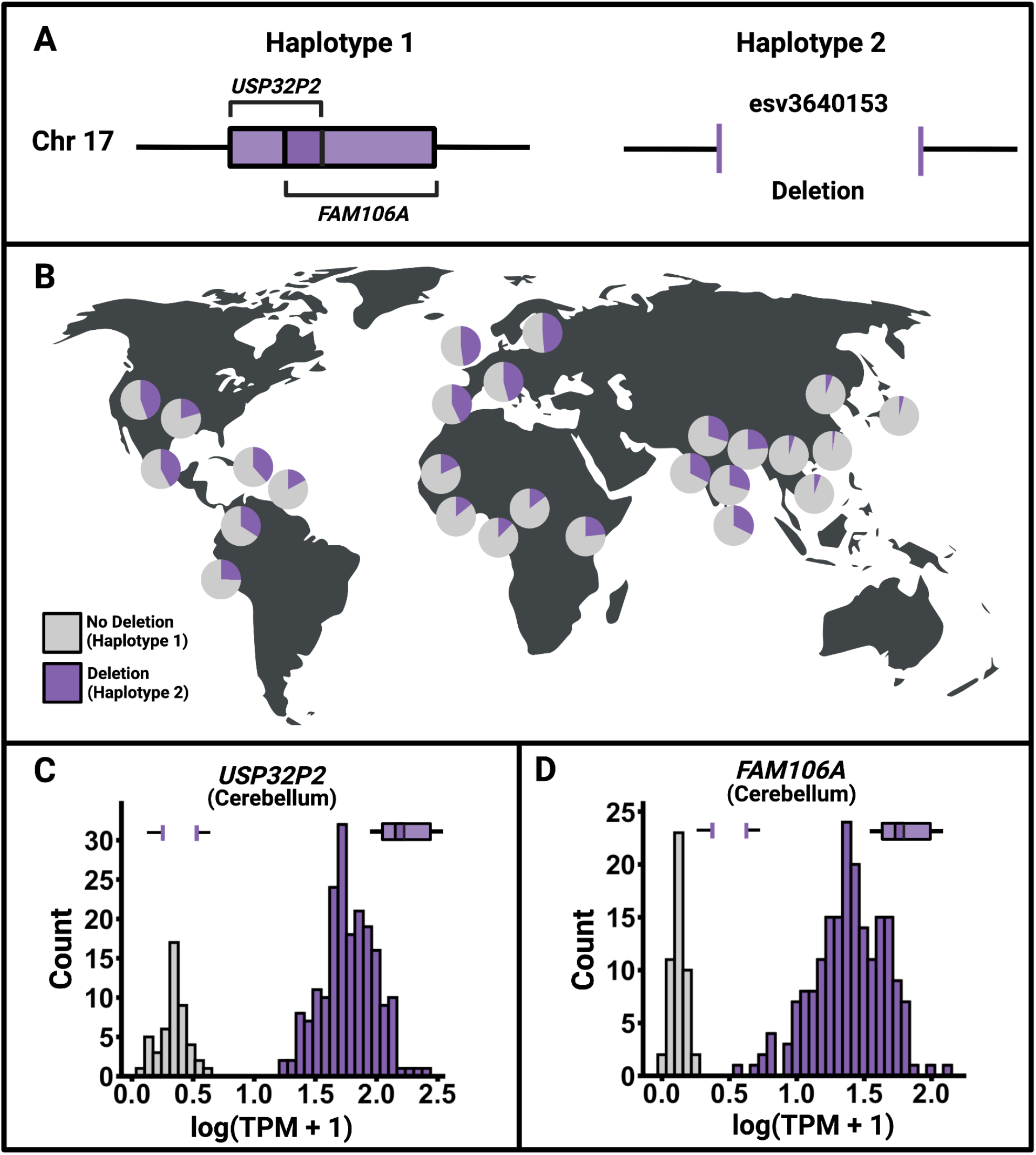
An example of a polymorphic gene deletion resulting in universally switch-like gene expression. **A**. *FAM106A* and *USP32P2* (not drawn to scale) are overlapping genes on chromosome 17. Two alternative haplotype classes exist for these genes: one in which both genes are completely deleted and the other without the deletion. **B**. Frequency distribution of the deletion across diverse populations. Each pie chart represents one of the 26 populations from the 1000 Genomes Project. Purple indicates the frequency of the deletion, while gray indicates the frequency of the alternative haplotype. **C-D**. Expression level distribution in the cerebellum (as an example) across individuals for *FAM106A* and *USP32P2*, respectively. The gene deletion presumably leads to the switched-off expression state in both genes.

We caution that we base our results regarding bimodality on expression at the RNA level. The bimodal expression of genes across individuals at the RNA level may not necessarily lead to bimodal expression at the protein level. For example, the universally switch-like expression of *RPS26* at the RNA level can be explained by a single nucleotide variant (rs1131017) in the gene’s 5’-untranslated region (UTR). In particular, *RPS26* has three transcription states based on the SNV genotypes. The ancestral homozygote C/C corresponds to a high transcription state, the heterozygote C/G to a medium state, and the derived homozygote G/G to a low state (See **Supplement** for a discussion on why an expression distribution driven by three genotypes at a polymorphic site might still appear bimodal). Remarkably, this pattern is reversed at the translation level ^24^: Messenger RNA carrying the derived G allele produces significantly more protein. This reversal may be due to a SNV in the 5’-UTR that can abolish a translation-initiation codon ^25^. This finding demonstrates how the same SNV can regulate a gene’s expression level in opposite directions during transcription and translation. This multi-level regulation in opposite directions likely serves to dampen protein expression variability. It has been shown previously that RNA variability is greater than protein variability in primates ^26,27^; the presence of dampening variants discussed here may be one reason behind these findings. Such compensatory mechanisms for gene expression remain fascinating areas for future research.

### Tissue-specific switch-like genes have a shared regulatory framework

Tissue-specific expression patterns are crucial for tissue function. Thus, we now turn our attention to tissue-specific switch-like genes. We found that the stomach, vagina, breast, and colon show a higher number of tissue-specific switch-like genes compared to other tissues (**Figure 5A**), after controlling for confounding factors (**Methods; Supplement; Table S6**). Furthermore, within these tissues, the expression of switch-like genes is not independent; instead, they exhibit high pairwise co-expression between genes (**Figure 5B-C; Table S7**). Hence, tissue-specific switch-like genes tend to be either all switched off or switched on within an individual. This result suggests a shared regulatory mechanism for the expression of these genes in each tissue. Given that hormonal regulation plays a substantial role in shaping tissue-specific expression patterns ^28,29^, we hypothesize that hormones may regulate genes that are bimodally expressed in specific tissues (cluster 1; **Figure 2B**).

**Figure 5.**
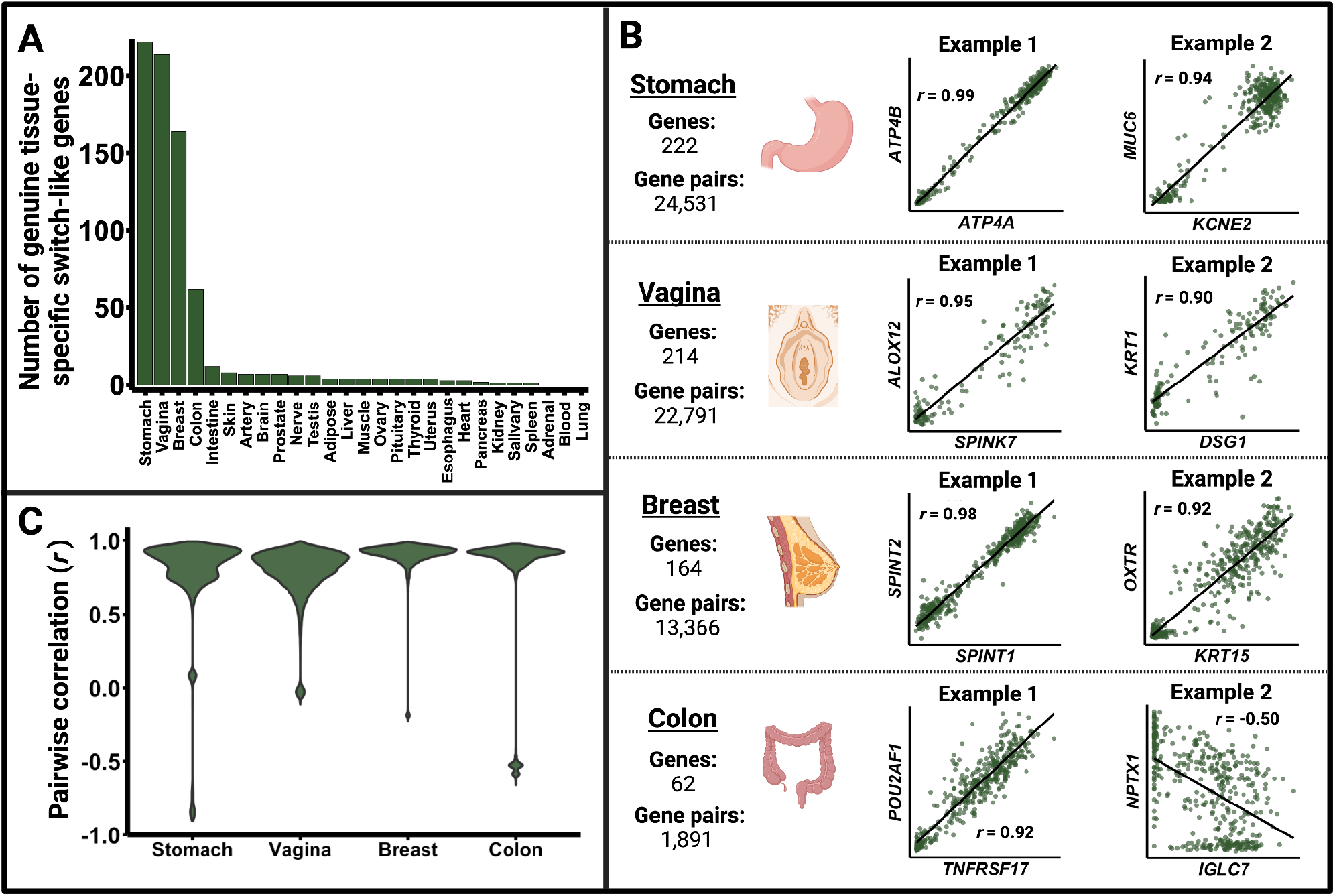
Characterization of genuine tissue-specific switch-like genes (cluster 1). The results shown here exclude genes that showed switch-like expression due to confounding factors like ischemic time. **A**. Number of tissue-specific switch-like genes showing bimodal expression in each of the 27 tissues. The stomach, vagina, breast, and colon show disproportionately more tissue-specific switch-like genes than other tissues. **B**. An illustration of how Pearson’s correlation coefficients were calculated for each pair of bimodally expressed tissue-specific switch-like genes within the stomach, vagina, breast, and colon. We show the scatterplots for two arbitrarily chosen gene pairs for each of the four tissues. The axes in each dot plot represent the log(TPM + 1) for the labeled gene in the relevant tissue. Panel C was generated using the pairwise correlation coefficients thus obtained. **C**. Tissue-specific switch-like genes within the four tissues shown are highly co-expressed. Tissue-specific master regulators, such as endocrinological signals, likely drive their concordant on and off states.

Sexual differences in hormonal activity are well documented ^30,31^. To explore this further, we investigated whether hormone-mediated sex-biased expression underlies the co-expression of tissue-specific switch-like genes within tissues. Under this scenario, a gene would be largely switched on in one sex and off in the other in a given tissue. Among tissue-specific switch-like genes, we identified 186 gene-tissue pairs with sex-biased bimodal expression (**Figure 6A; Table S8**). These instances are biologically relevant; for example, we found switch-like immunoglobulin genes with female-biased expression in the thyroid, heart, tibial nerve, and subcutaneous adipose tissue. This observation may relate to previous findings ^32,33^ of higher antibody responses to diverse antigens in females than in males.

**Figure 6.**
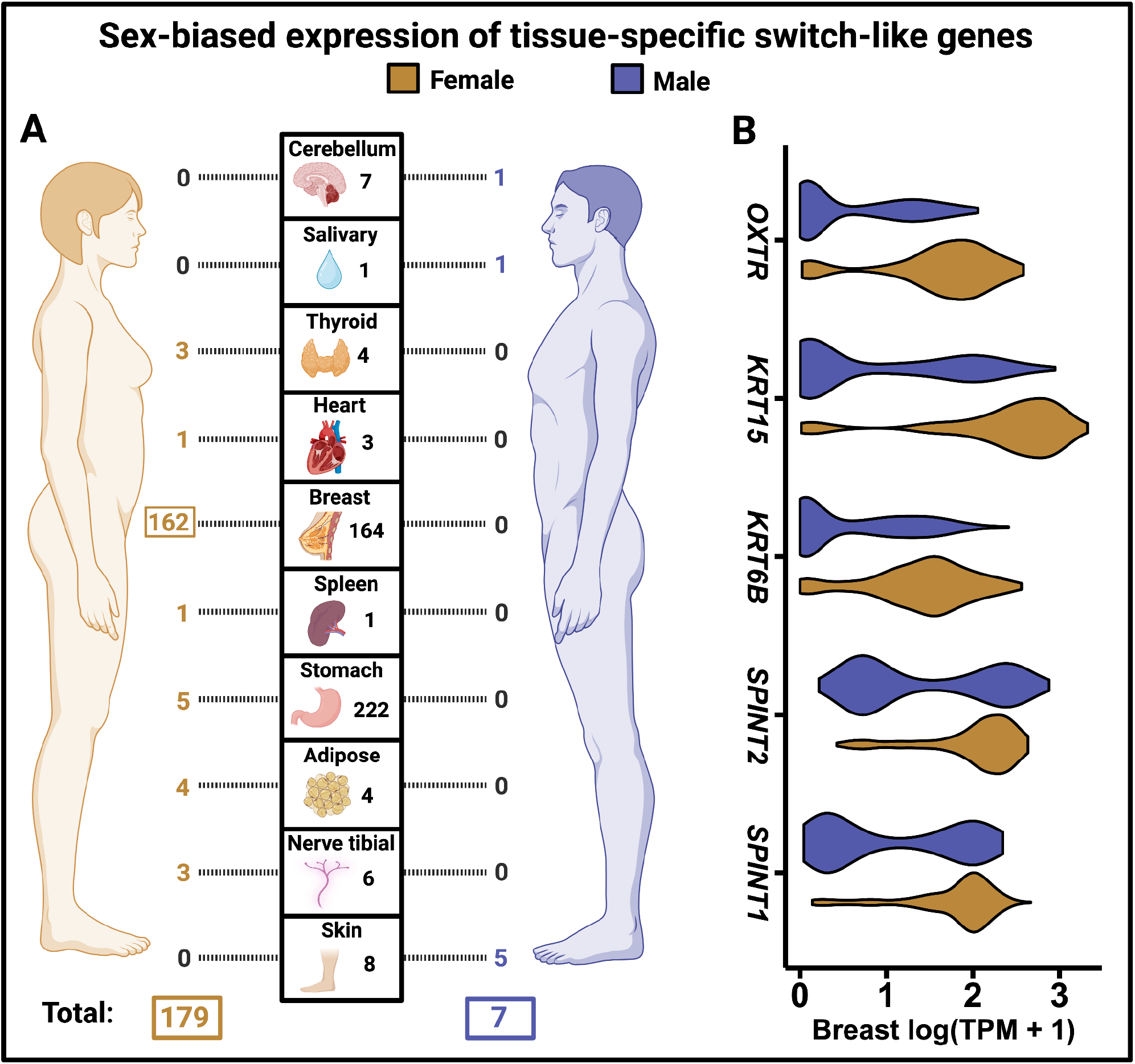
Sex-biased expression of tissue-specific switch-like genes (cluster 1). **A**. Number of tissue-specific switch-like genes that show female- and male-biased expression. Only those tissues are shown that have at least one tissue-specific switch-like gene showing sex bias. The number in the central grid next to each tissue image represents the number of genuine tissue-specific switch-like genes in that tissue. In orange, the numbers to the left of the central grid indicate the count of female-biased genes in each of the 10 tissues shown. In blue, the numbers to the right of the grid indicate the count of male-biased genes. **B**. Violin plots showing the expression level distribution in the breast for five female-biased tissue-specific switch-like genes discussed in the main text.

More dramatically, we found that 162 out of 164 tissue-specific switch-like genes (cluster 1) in the breast tissue are female-biased, explaining their correlated expression levels (**Figure 6A**). However, the sex-based disparity in the on-versus-off states of these genes is not absolute, but rather a statistical tendency. In other words, the gene is not switched off in all males and switched on in all females. Instead, the proportion of individuals with the gene switched on significantly differs between sexes. Notably, multiple sex-biased switch-like genes—including *SPINT1* and *SPINT2* ^34^, multiple keratin genes ^35^, and the oxytocin receptor gene ^36,37^ (*OXTR;* **Figure 6B**)—in the breast tissue are differentially expressed in breast cancers relative to matched non-cancerous tissues. Future investigations could reveal whether the toggling of these genetic switches affects breast cancer risk in females. We caution that sex-biased switch-like expression in the breast may result from differences in cell-type abundance between females and males. Nevertheless, the differential expression of some genes between sexes might developmentally drive such differences in cell-type abundance. In summary, our results indicate that sex is a major contributor to bimodal gene expression, with breast tissue standing out as particularly sex-biased in this context.

We note that the intra-tissue co-expression of tissue-specific switch-like genes in the stomach and colon cannot be explained by sex. By biological definition, the variation in vaginal expression levels in our sample is not sex-biased. Thus, the intra-tissue co-expression of tissue-specific switch-like genes in the stomach, colon, and vagina may be explained by one of two reasons: 1) Most of the tissue-specific switch-like genes in each tissue are directly regulated by the same hormone in that tissue, or 2) Most of the tissue-specific switch-like genes in each tissue are regulated by the same transcription factor which is, in turn, under regulation by a hormone or other cellular environmental factors. In the case of hormonally controlled gene expression, genes are likely switched off when the systemic hormone levels drop below a certain threshold. We will discuss this idea further, specifically for the vagina, later in the text.

### Concordantly switched-off genes in the stomach may indicate a predisposition to gastric cancer

Gene expression levels have been studied as a diagnostic marker for disease states ^38^. Therefore, we asked whether tissue-specific switch-like genes co-expressed with each other across individuals are linked to human disease, with each of the two expression states corresponding to different risks. To address this question, we investigated whether the identified switch-like genes in a given tissue are overrepresented among genes implicated in diseases of the same tissue.

We overlapped the switch-like genes in the stomach with a previously published list ^39^ of differentially expressed genes in gastric carcinomas. We found that switch-like genes in the stomach are significantly enriched (41-fold enrichment, *p*<10^-4^) among genes that are downregulated in gastric carcinomas. Specifically, nine switch-like genes are downregulated in gastric carcinomas (*ATP4A, ATP4B, CHIA, CXCL17, FBP2, KCNE2, MUC6, TMEM184A*, and *PGA3*). Additionally, these nine genes are concordantly expressed in 92.5% (332/359) of the stomach samples, being either all switched off or on in a given individual (**Methods**). Our data suggest that individuals with these nine genes switched off in the stomach may be susceptible to developing cancers. This preliminary observation provides exciting avenues to investigate both the cause of the concordant toggling of these genes and their potential role in cancer development.

### Concordantly switched-off genes result in vaginal atrophy

We found that switch-like genes in the vagina are significantly overrepresented (44-fold enrichment; *p*<10^-4^; see methods) among genes linked to vaginal atrophy in postmenopausal women. Vaginal atrophy, affecting nearly half of postmenopausal women, is triggered by sustained low levels of systemic estrogen and is marked by increased microbial diversity, higher pH, and thinning of the epithelial layer in the vagina ^40,41^. It is also known as atrophic vaginitis, vulvovaginal atrophy, estrogen-deficient vaginitis, urogenital atrophy, or genitourinary syndrome of menopause, depending on the specialty of the researchers. Symptoms experienced by women include dryness, soreness, burning, decreased arousal, pain during intercourse, and incontinence ^42^. Our analysis of switch-like genes in the vagina provides new insights into the development of vaginal atrophy.

Specifically, we overlapped a previously published list ^43^ of genes that are transcriptionally downregulated in vaginal atrophy with our list of bimodally expressed genes in the vagina. We found that the genes *SPINK7, ALOX12, DSG1, KRTDAP, KRT1*, and *CRISP3* are both bimodally expressed in the vagina and transcriptionally downregulated (presumably switched off) in women with vaginal atrophy **(Figure 7A)**. We refer to these genes as “atrophy-linked switch-like genes.” Indeed, these six genes are either all switched on, or all switched off concordantly in 84% (131/156) of the vaginal samples we studied. The pairwise concordance rates (percentage of individuals with both genes switched on or both genes switched off) for these genes are shown in **Figure 7B**. Among postmenopausal women with this concordant gene expression, 50% are in the “off” state – a fraction that closely matches the prevalence of vaginal atrophy in postmenopausal women ^40,44^. Therefore, our data suggest that estrogen-dependent transcription underlies concordant expression of atrophy-linked switch-like genes, with the “off” state of these genes associated with vaginal atrophy.

**Figure 7.**
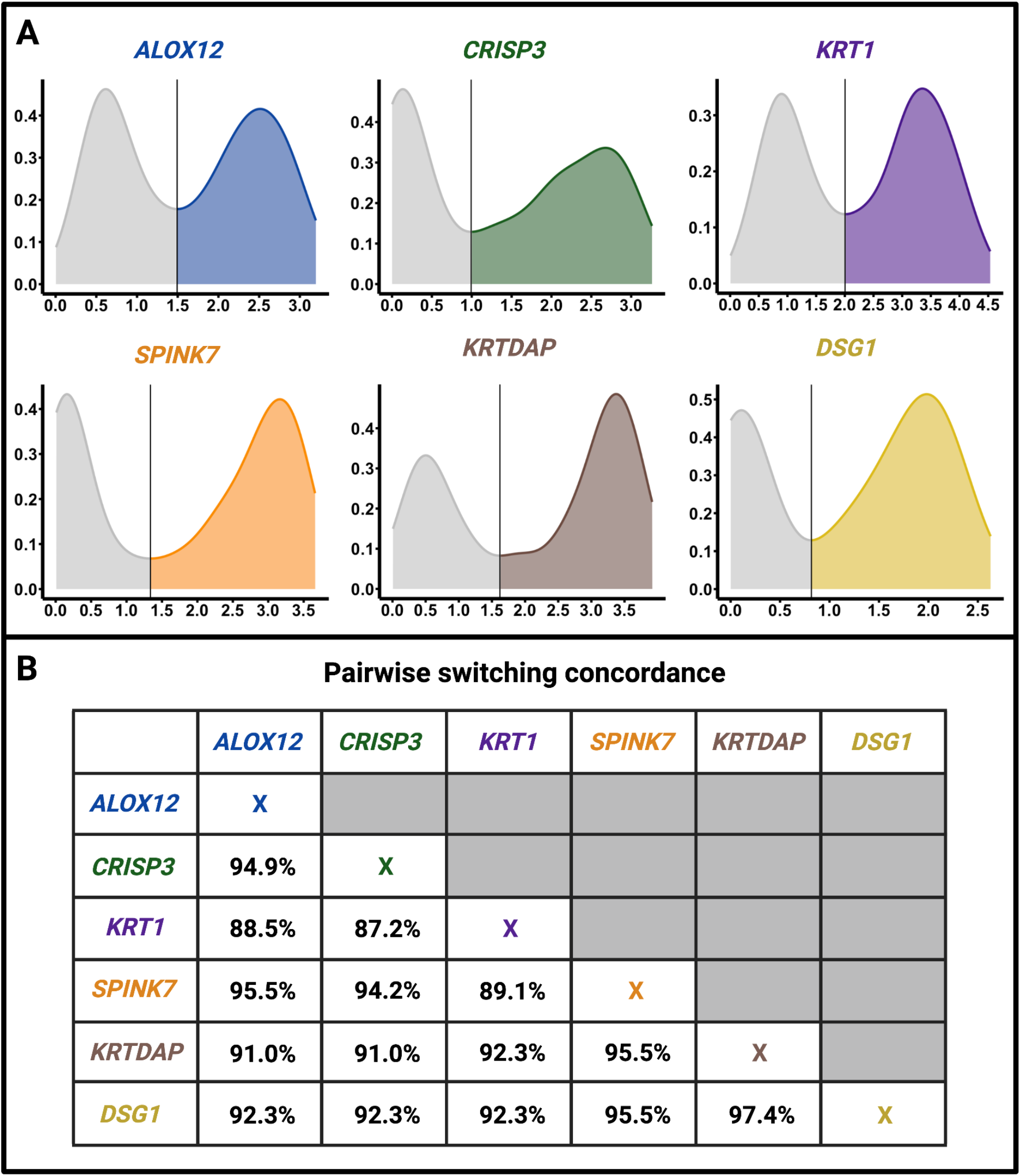
Atrophy-linked switch-like genes tend to be either all switched off, or all switched on within individuals. **A**. The distribution of expression levels in the vagina of the six switch-like genes implicated in vaginal atrophy. The x-axes represent log(TPM +1) values for each gene in the vagina, and the y-axes represent the probability density. We obtained the probability densities using kernel density estimation. In each case, the global minimum (excluding endpoints) is considered the switching threshold. A gene is deemed “on” in an individual if the expression level is above this threshold; otherwise, the gene is deemed “off.” **B**. Pairwise concordance rates (percentage of individuals in which the two genes are either both switched on or both switched off).

For background, the vaginal epithelial layers are differentiated from the inside out. The basal and parabasal layers of the epithelium consist of mitotic progenitor cells with differentiation potential, while the outermost layer comprises the most differentiated cells ^45,46^. When basal and parabasal cells stop proliferating, the death of mature cells leads to a thin epithelium, and the symptoms of vaginal atrophy appear. Given this background, atrophy-linked switch-like genes may either be a cause or a consequence of vaginal atrophy. In particular, if an atrophy-linked switch-like gene encodes a protein necessary for the continued proliferation and differentiation of basal and parabasal cells, we call it a “driver” gene. In the absence of the driver gene’s protein, cell differentiation ceases, and the outer layer gradually disappears, resulting in vaginal atrophy (**Figure 8A**). On the other hand, if the product of an atrophy-linked switch-like gene is not required for basal and parabasal cell proliferation, we refer to it as a “passenger” gene, borrowing the terminology from cancer literature ^47^. In healthy vaginas with a thick epithelium, there are more cells in which passenger genes would be expressed. By contrast, in atrophic vaginas, the epithelium thins, resulting in fewer cells where these genes can be expressed. This contrast would lead to the bimodal expression of passenger genes across vagina samples in whole-tissue RNA-sequencing datasets. We hypothesize that at least some of the atrophy-linked switch-like genes are driver genes.

**Figure 8.**
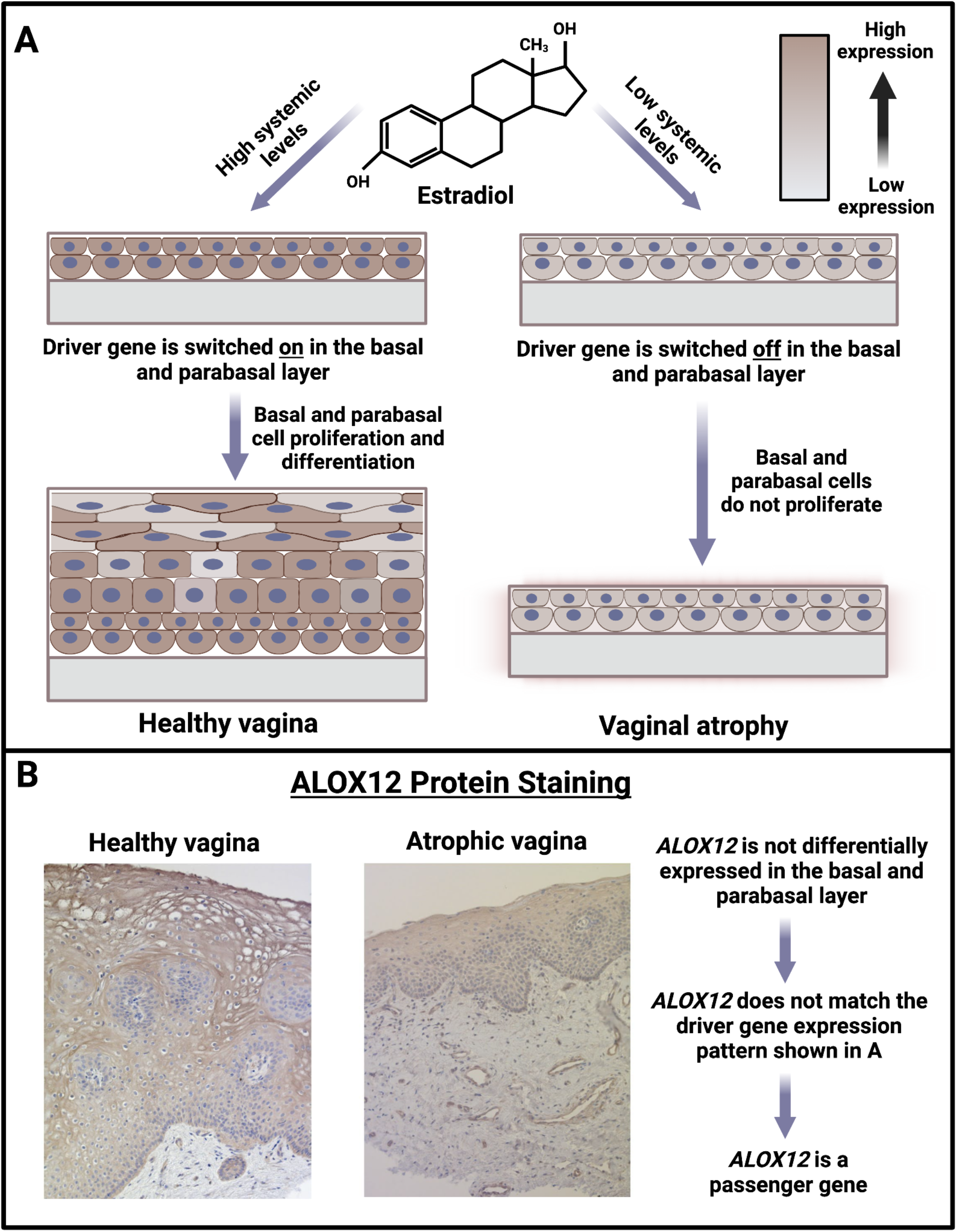
*ALOX12* is a passenger gene. **A**. Model for the etiology of vaginal atrophy. High levels of estrogen keep the driver genes switched on in basal and parabasal epithelium, impelling basal and parabasal cells to proliferate and mature, resulting in healthy vaginal mucosa. Conversely, low levels of estrogen switch off the driver genes. The lack of basal and parabasal cell proliferation leads to a thin vaginal epithelium, resulting in vaginal atrophy. **B**. Representative immunohistochemical staining of Arachidonate 12-Lipoxygenase (ALOX12) in vaginal tissue. We show healthy vaginal tissue from a woman with higher systemic estrogen levels and a thicker vaginal epithelial layer, along with atrophic vaginal tissue from a woman with low systemic estrogen levels and a thinner vaginal epithelial layer. There is no difference in *ALOX12* expression in the basal or parabasal cells between healthy and atrophic epithelium, implicating it as a passenger gene. Images taken with Axio Observer Z1 (Carl Zeiss AG) with a 40X objective.

Two key findings allowed us to construct this hypothesis. Firstly, switch-like genes in the vagina show a 26-fold ontological enrichment for the establishment of the skin barrier (FDR=1.26 x10^-6^) and a 25-fold enrichment for keratinocyte proliferation (FDR=1.75 x 10^-4^), both related to epithelial thickness and differentiation. Notably, two atrophy-linked switch-like genes in the vagina that we identified, *KRTDAP* and *KRT1*, are crucial for the differentiation of epithelial cells in the vagina ^48,49^. Protein stainings available through Human Protein Atlas ^50^ show that all six atrophy-linked switch-like genes are expressed at the protein level, predominantly in the vaginal epithelium. Secondly, administering 17β-estradiol (a type of estrogen) to postmenopausal women with vaginal atrophy leads to the upregulation of the same six genes, causing symptoms to subside ^51^. According to our hypothesis, administering estrogen activates the expression of the driver switch-like genes in the vagina, resuming the proliferation of basal and parabasal cells in the epithelium. This process leads to the reformation of a thick and healthy vaginal mucosa, thereby alleviating the symptoms of vaginal atrophy.

Thus, it is essential to distinguish driver genes from passenger genes to understand the etiology of vaginal atrophy. However, we expect driver and passenger genes to show the same expression patterns in healthy versus atrophic vaginas using bulk RNA-sequencing data. In order to make this distinction, we need comparative expression data, specifically from the basal and parabasal epithelium from healthy versus atrophic vaginas. We expect driver genes to be differentially expressed in the basal and parabasal layers of the epithelium. By contrast, we expect passenger genes to show no differential expression in the basal and parabasal layers between healthy and atrophic vaginas.

To look at the expression levels in the basal and parabasal layers of the epithelium, we arbitrarily chose *ALOX12* from the six atrophy-linked switch-like genes for immunohistochemical staining of its protein product in the vaginal mucosa (which includes the epithelium and the underlying connective tissue). We found that the ALOX12 protein is present in the epithelial cells, and its abundance directly correlates with epithelial thickness, as expected from our RNA-sequencing results. However, we found no significant difference in the staining of the ALOX12 protein in the basal or parabasal epithelial layers between healthy and atrophic samples **(Figure 8B)**. This suggests that the gene is not differentially expressed in the basal or parabasal layers of the vaginal epithelium between healthy and atrophic vaginas. Therefore, *ALOX12* is a passenger gene for vaginal atrophy. Comparative immunohistochemical staining of the protein product of the other five atrophy-linked switch-like genes may identify the driver gene in the future. Indeed, the KRT1 protein is recognized as a marker of basal cell differentiation in mouse vaginas ^52^, a finding that may also be true for humans. Overall, our results open up several new paths for potential pre-menopausal risk assessment and intervention frameworks targeting cell differentiation pathways in the clinical setting.

## Discussion

In this study, we investigated factors underlying switch-like gene expression and its functional consequences. Our systematic analysis revealed 1,013 switch-like genes across 943 individuals. Some of these genes show bimodal expression across individuals in all tissues, suggesting a genetic basis for their universally switch-like behavior. We found several single nucleotide and structural variants to explain the switch-like expression of these genes. Most of the switch-like genes, however, exhibit tissue-specific bimodal expression. These genes tend to be concordantly switched on or off in individuals within the breast, colon, stomach, and vagina. This concordant tissue-specific switch-like expression in individuals is likely due to tissue-specific master regulators, such as endocrinological signals. For example, in the vagina, switch-like genes tend to get concordantly switched off in a given individual when systemic estrogen levels fall below a certain threshold. On the biomedical front, our work linked switch-like expression to the susceptibility to gastric cancer and vaginal atrophy. Furthermore, this study has paved two major paths forward toward early medical interventions, as discussed below.

First, we emphasize that bimodal expression that is correlated across all tissues is driven by genetic polymorphisms. However, the genetic bases for 13/23 universally switch-like genes remain elusive. We propose that the underlying genetic bases for these universally switch-like genes are structural variants, which are not easily captured by short-read DNA sequencing. These structural variants may be discovered in the future as population-level long-read sequencing becomes more common. The first biomedical path forward is to use long-read DNA sequencing to pinpoint the genetic polymorphisms responsible for the bimodal expression of disease-related genes. Of particular interest are the genes *CYP4F24P* and *GPX1P1*, both long non-coding RNAs, which are implicated in nasopharyngeal cancer. The genetic basis for their bimodal expression remains unknown. *CYP4F24P* is significantly downregulated in nasopharyngeal cancer tissues ^53^, while *GPX1P1* is significantly upregulated in nasopharyngeal carcinomas treated with the potential anticancer drug THZ1 ^54^. Investigating whether individuals with naturally switched-off *GPX1P1* and *CYP4F24P* are at a higher risk of nasopharyngeal cancer will enable genotyping to identify individuals at elevated risk for nasopharyngeal cancer, facilitating early interventions and improving patient outcomes.

Secondly, switch-like genes present a promising avenue for exploring gene-environment interactions, an area of growing interest. Recent studies indicate that environmental factors can significantly modulate genetic associations ^55,56^. Polymorphisms that result in switch-like gene expression have already been linked to several diseases within specific environmental contexts ^57^. For instance, the deletion of *GSTM1* has been associated with an increased risk of childhood asthma, but only in cases where the mother smoked during pregnancy ^58^. Even more critically, switch-like genes potentially create unique cellular environments that could modulate the impact of genetic variations. We hypothesize that switch-like expression can produce diverse cellular environments, whether in a single gene (as in genetically determined cases) or in multiple genes (as in tissue-specific, hormonally regulated cases). These environments may, in turn, influence the effect of genetic variations and their associations with disease. Thus, much like current gene-environment association studies that control for factors such as birthplace, geography, and behaviors like smoking, it is conceivable that controlling for switch-like gene expression states could enhance the power of such studies. By cataloging these switch-like genes and developing a framework to classify them as “on” or “off” in various samples, our work lays the groundwork for more robust association studies in future research.

In summary, our study has significant implications for understanding the fundamental biology of gene expression regulation and the biomedical impact of switch-like genes. Specifically, it contributes to the growing repertoire of methods for determining individual susceptibility to diseases, facilitating early therapeutic interventions. By providing a new approach to studying gene expression states, our study will enhance the predictive accuracy of disease susceptibility and improve patient outcomes.

## Acknowledgment

O.G. and N.M. acknowledge support from the National Institute of General Medical Sciences (under grant no.1R01GM148973-01). N.M. also acknowledges support from the Japan Science and Technology Agency (JST) Moonshot R&D (under grant no.JPMJMS2021), the National Science Foundation (under grant no.2052720), and JSPS KAKENHI (under grant no.JP 24K14840). O.G. acknowledges support from the National Science Foundation (under grant nos.2049947 and 2123284). The funders had no role in study design, data collection and analysis, decision to publish, or preparation of the manuscript.

## METHODS

### Data

The Genotype-Tissue Expression (GTEx) project is an ongoing effort to build a comprehensive public resource to study tissue-specific gene expression and regulation. The data we use are transcript per million (TPM) obtained from human samples across 54 tissues and 56,200 genes (as of December 1st, 2023). We excluded laboratory-grown cell lines from our analysis. Since we need a reasonable number of individuals from each tissue, we excluded tissues with less than 50 individuals for our calculations. Of the remaining tissues, there were instances of multiple tissues from the same organ. In such cases, we randomly chose one tissue per organ. We thus focus our analysis on 27 tissues (**Figure 1**). Additionally, we retained only those genes for which the mean TPM across individuals was greater than 10 in at least one of the 27 focal tissues. This filter was applied because the analysis of lowly expressed genes may lead to false positive calls for bimodal expression and, as a result, to assign biological significance to cases where there is none. After these filtering steps, we are left with TPM data from 19,132 genes in each of the 27 tissues. We note that each tissue contains data from a different number of samples (individuals), totaling 943 across tissues. We will refer to this set of 19,132 genes as *G* in our equations and the rest of the methods.

### Dip test

There are many tests of bimodality of gene expressions ^16,59^. We use a dip test described as follows. We denote by *S*_*i*_ the number of samples (individuals) available for tissue *i*. We also denote by *x*_*g*,*i*,*s*_ the TPM value for gene *g* in tissue *i*, for sample *s* ∈ {1,…, *S*_*i*_} and *g* ∈ *G*. According to convention, we log-transform the TPM, specifically by log(*x*_*g*,*i*,*s*_+1) ^60^ to suppress the effect of outliers; TPM is extremely large for some samples. Note that log(*x*_*g*,*i*,*s*_+ 1) conveniently maps *x*_*g*,*i*,*s*_= 0 to 0. For each pair of gene *g* and tissue *i*, we carried out a dip test, which is a statistical test for multimodality of distributions, on the distribution of log(*x*_*g*,*i*,*s*_+ 1) across the samples *S*_*i*_. We performed the dip test using the dip.test() function within the “diptest” package in R, with the number of bootstrap samples equal to 5000. We applied the Benjimini-Hochberg procedure for multiple hypothesis correction to the results with a false discovery rate of 5%. Additionally, to reduce false positive calls of bimodal expression, we only retained results where the dip statistic 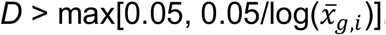, where

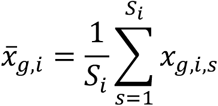

We obtained this threshold of 0.05 by visual inspection of log(*x*_*g*,*i*,*s*_+ 1) distributions in the stomach and adipose subcutaneous tissues, starting with those with the highest values of *D*. For statistically significant results, the distribution was almost always bimodal if *D* exceeded 0.05. The only exceptions were genes with low 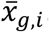. Thus, we penalized gene-tissue pairs with low 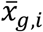 across samples by requiring a higher *D* in order to classify them as bimodally distributed. Genes identified as bimodally distributed in at least one tissue are referred to as “switch-like” genes.

### Tissue-to-tissue co-expression of genes

We sought to identify switch-like genes whose expression exhibits bimodal expression in all tissues. One seemingly straightforward approach is to count the number of tissues showing bimodal distribution of expression levels for each gene. However, even if a gene genuinely exhibits bimodal expression across all tissues, our methodology may fail to recognize it as such if the mean expression levels 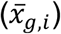 of the gene are low in some tissues. This is because our effect size threshold penalizes gene-tissue pairs with low 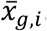. Moreover, if gene expression follows a bimodal distribution across all tissues, then it does so likely due to a genetic polymorphism affecting expression. Thus, the expression of such genes would be highly correlated between pairs of tissues. Given this insight, discovering universally bimodal genes is more tractable using tissue-to-tissue co-expression of each gene.

For each gene, we construct the co-expression matrix among pairs of tissues as follows. To calculate the co-expression between a pair of tissues, we need to use the samples whose TPM is measured for both tissues ^61^. In general, even if the number of samples is large for both of the two tissues, it does not imply that there are sufficiently many common samples. Therefore, using the sample information described in GTEx_Analysis_v8_Annotations_SampleAttributesDD.xlsx in the GTEx data portal, we counted the number of samples shared by each tissue pair and excluded the 41 tissue pairs that share less than 40 samples. For each of the remaining 27 x 26/2 - 41 = 310 tissue pairs, we denote by *S*_*i*,*j*_ the number of samples shared by the two tissues *i* and *j*. We also denote by *x*_*g*,*i*,*s*_ and *x*_*g*,*j*,*s*_ the TPM value for gene *g* in tissues *i* and *j*, respectively, for sample *s* ∈ {1, 2, … , *S*_*i, j*_}. Then, we calculated the Pearson correlation coefficient between log(*x*_*g*,*i*,*s*_ + 1) and log(*x*_*g*,*j*,*s*_ + 1) across the *S*_*i*,*j*_ samples and used it as the strength of the co-expression of gene *g* between tissues *i* and *j*. Specifically, we calculate

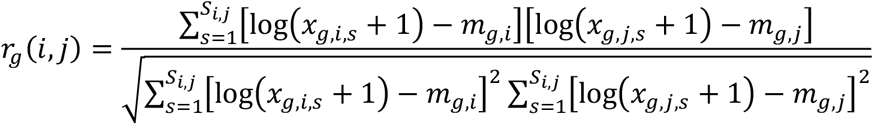

where

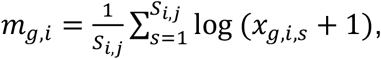

and

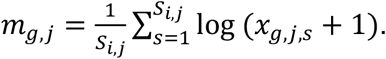

For each gene *g*, we then vectorize the correlation matrix, (*r*_*g*_(*i, j*)), into a 310-dimensional vector. If, for a given gene, *g*, log(*x*_*g*,*i*,*s*_+ 1) or log(*x*_*g*,*j*,*s*_+ 1) were 0 across all *S*_*i*,*j*_ samples for any of the 310 tissue pairs, the gene was removed. In this process, 28 out of 1,013 switch-like genes were removed. Note that the correlation matrix is symmetric, so we only vectorize the upper diagonal part of the matrix. We denote the generated vector by 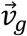. Vector 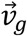 characterizes the gene. We ran a principal component analysis (PCA), using the prcomp() function in R, on vectors, 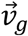 for all genes for which we could calculate *r*_*g*_(*i, j*) for all 310 tissue pairs. In parallel, we also ran PCA on only the set of vectors (genes) characterizing only the 985 (1013 - 28) switch-like genes.

In the space spanned by the first two principal components, we calculated the pairwise distance between genes using the dist() function in R with method = “euclidean”. We then performed hierarchical clustering using the hclust() function with method = “complete”. Finally, we used the cuttree() function with *k*=2 and *k*=3 to obtain two and three clusters, respectively.

### Identifying the genetic basis of universal bimodality

In order to identify the genetic basis of bimodality for switch-like genes in cluster 2A, we obtained the coordinates of the genes for both hg19 and hg38 using their Ensembl IDs as keys through Ensembl BioMart. We obtained coordinates of common structural variants using both the 1000 genomes project (hg19) ^62^ and the HGSV2 dataset (hg38) ^63^. We performed an overlap analysis using BedTools ^64^ to identify polymorphic deletions of or insertions into these genes. We thus obtained five universally bimodal genes being affected by structural variants. These were *USP32P2, FAM106A, GSTM1, RP11-356C4*.*5*, and *CYP4F24P*. Additionally, we obtained the GTEx dataset for the expression quantitative trait loci (eQTL). We identified genes in cluster 2A that had at least one eQTL, which was consistently associated with either increased or decreased expression of a given gene across all 27 tissues analyzed. We thus obtained five genes from cluster 2A whose expression was associated with a short variant across tissues. These were *NPIPA5, RPS26, PSPHP1, PKD1P2*, and *PKD1P5*.

### Controlling for confounders

A bimodal distribution of expression levels of universally switch-like genes is unlikely to be driven by confounding factors such as ischemic time, and time spent by the tissue in chemical fixatives (PAXgene fixative). For example, the expression of genes on the male-specific region of chromosome Y is bimodally distributed across tissues regardless of confounding factors because females do not possess these genes. Similarly, regardless of confounding factors, *USP32P2* is bimodally distributed due to a polymorphic gene deletion. However, tissue-specific switch-like genes are particularly prone to being affected by confounding variables. Specifically, we investigated whether the switch-like expression of genes can be explained by ischemic time and PAXgene fixative using the following approach.

Ischemic time for a sample *s* in a given tissue *i*, denoted by *k*_*i*,*s*_, is a continuous variable representing the time interval between death and tissue stabilization. Time spent by a tissue *i* from a sample *s* in PAXgene fixative, denoted by *f*_*i*,*s*_, is also a continuous variable. For each gene-tissue pair (*g, i*), we calculated, across the *S*_*i*_ samples, the Pearson correlation between 1) log(1 + *x*_*g*,*i*,*s*_) and *k*_*i*,*s*_ and 2) log(1 + *x*_*g*,*i*,*s*_) and *f*_*i*,*s*_. For each tissue *i* and confounder *c*, where *c* ∈ {*k*_*i*,*s*_, *f*_*i*,*s*_}, we denote the correlation coefficient between log(1 + *x*_*g*,*i*,*s*_) and *c* as *r*_*g*,*i*,*c*_.

We partition the set of switch-like genes into two subsets: cluster 1 and cluster 2 (the union of clusters 2A and 2B). We treat cluster-2 genes as internal controls since their correlated bimodal expression across tissues is robust to the presence of confounding factors. Thus, we eliminated a cluster-1 gene *g*1 if, for any confounder 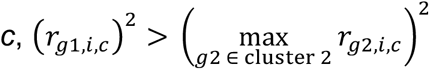.

### Gene-to-gene co-expression within tissues

We performed gene-to-gene co-expression analysis within the stomach, breast, vagina, and colon tissues. In a given tissue *i*, we denote the set of genuine cluster-1 genes (excluding genes affected by confounding variables) by *C*_*i*_. Then, for *i* ∈ {stomach, breast, vagina, colon}, we calculated the Pearson correlation, across the *S*_*i*_ samples, between log(*x*_*g*,*i*,*s*_ + 1) and log(*x*_*h*,*i*,*s*_ + 1) for every *g, h* ∈ *C*_*i*_ where *g* ≠ *h*.

### Quantifying sex bias in cluster-1 gene expression

For every gene-tissue pair (*g*,*i*), where *g* is a switch-like gene, and *i* is a tissue common to both sexes, we tested the hypothesis that the distribution of log(*x*_*g*,*i*,*s*_+ 1) across male samples differed from that across female samples using the Wilcoxon rank-sum test.

We applied the Benjamini-Hochberg procedure of multiple hypotheses correction with FDR = 5*s*. We quantified the effect size of the sex bias using Cohen’s *d*. Statistically significant results were considered to represent true sex bias only if |*d*| > 0.2 ^65^.

### Enrichment of switch-like genes among disease-linked genes

We performed enrichment analysis for switch-like genes in the stomach and vagina that are downregulated in gastric cancer and vaginal atrophy, respectively. We denote the set of genes downregulated in disease *y* as *Z*_*y*_, where *y* ∈ {gastric cancer, vaginal atrophy}. We calculated the fold enrichment of genuine cluster-1 genes in the stomach among genes downregulated in gastric cancer by:

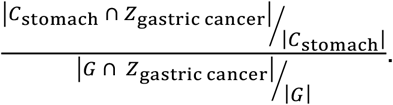

We calculated the fold enrichment of genuine cluster-1 genes in the vagina among genes downregulated in vaginal atrophy by:

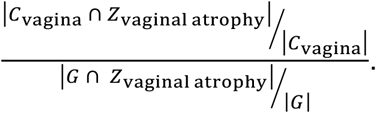

To calculate the *p*-values associated with these enrichments, we obtained 10,000 uniformly random samples (with replacement) of size |*C*_*i*_| from *G*. The *p-*value for the enrichment of switch-like genes in tissue *i* among genes linked to disease *y* is then given by the fraction of random samples among the 10,000 samples for which |*q*_*j*_ ∩ *Z*_*y*_| > |*C*_*i*_ ∩ *Z*_*y*_|. Here, *q*_*j*_ is the set of genes in random sample *j* where *j* ∈ {1, …, 10000}.

### Discretizing expression levels

We performed kernel density estimation using the density() function in R on the distributions of 1) log(*x*_*g*,stomach,*s*_ + 1) across the *S*_stomach_ samples for *g* ∈ *C*_stomach_ ∩ *Z*_gastric cancer_ ; and 2) log(*x*_*g*,vegina,*s*_+ 1) across the *S*_vegina_ samples for *g* ∈ *C*_vegina_ ∩ *Z*_veginal atrophy_.

We used the minimum of the estimated density as the switching threshold; if an individual had an expression level above the threshold in a given tissue, the gene was considered “on” in the individual in that tissue. The gene was considered “off” otherwise. We then calculate the concordance of expression among genes in any arbitrary set of switch-like genes *G*^*A*^ in a given tissue *i* as follows:

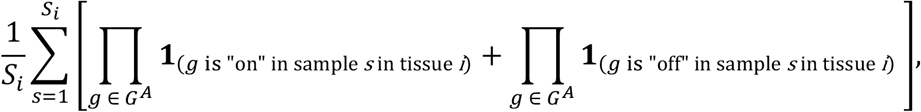

where **1**_(⋅)_ is the indicator function.

### Gene ontology enrichment of tissue-specific switch-like genes in the vagina

We performed Gene Ontology (GO) enrichment analysis for genes in *C*_vegina_ using the online database available at https://geneontology.org/ ^66^.

### Immunohistochemistry

Vaginal biopsies were taken by use of punch biopsies from postmenopausal women, fixed and stained as previously described by use of ALOX12 (HPA010691 polyclonal antirabbit, Sigma-Aldrich) ^67,68^.

## Supplementary Information

### Principal component analysis on tissue-to-tissue co-expression vectors

We applied a principal component analysis to the 19,132 vectors of tissue-to-tissue co-expression, one vector for each gene. We find that PC1 (**Figure 2A**), explaining 35.3% of the variation, is nearly perfectly correlated with mean tissue-to-tissue co-expression across tissue-tissue pairs (*r*^2^ = 0.998, *p*-value < 2.2×10^-16^; **Figure S1**). This result indicates that the 35.3% of the variation in the tissue-to-tissue co-expression of genes is primarily explained by the mean tissue-to-tissue co-expression of genes.

**Figure S1.**
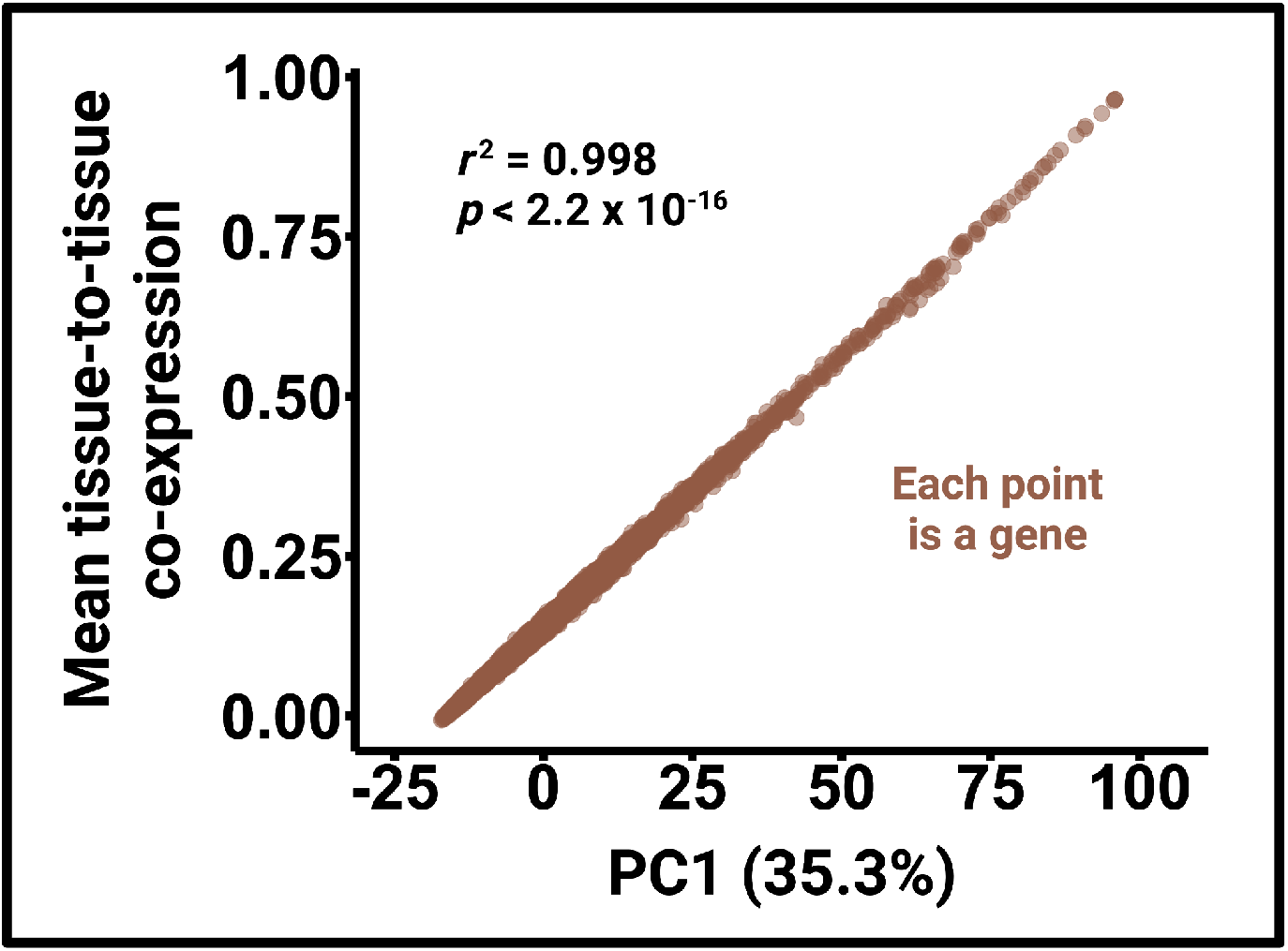
The mean tissue-to-tissue co-expression of genes shows a near-perfect correlation with PC1.

### Universally switch-like genes and their biomedical implications

In the main text, we discussed the *USP32P2* and *FAM106A*. Here, we discuss some other interesting examples of universally switch-like genes. The violin plots for the expression level distributions for all cluster-2A and cluster-2B switch-like genes not shown in the main text are present in **Figure S2** and **Figure S3**, respectively.

Firstly, a common ∼20kb whole-gene deletion (esv3587154) of the *GSTM1* gene ^69,70^ is associated with bladder cancer in humans ^71^. *GSTM1* is bimodally expressed across individuals in all tissues (**Figure S2D**) that we analyzed, as well as across multiple tumor types ^15^, with different expression peaks corresponding to differential prognoses among patients. These findings suggest a compelling hypothesis: the common deletion of *GSTM1*, maintained either by drift or balancing selection ^72^, has no significant effect on the health of non-cancerous individuals; however, it could have significant implications for prognosis once certain types of tumors develop. Therefore, screening patients with certain tumor types for the *GSTM1* deletion could significantly advance our ability to predict the course of tumor progression in an individualized manner.

Secondly, genes that are bimodally expressed across multiple tissues raise an evolutionary paradox. Typically, genes with a wide expression breadth (i.e., expression across a large number of tissues) affect fitness and are thus constrained at both the sequence and expression level ^26,73–75^. However, universally switch-like genes, despite having a high expression breadth, are not conserved at the expression level. This could imply different health consequences for individuals with off versus on state of the genes. For example, the universally switch-like gene RP4-765C7.2 (ENSG00000213058; **Figure S2K**) is upregulated in the peripheral blood mononuclear cells of patients with ankylosing spondylitis ^76^, eutopic endometrium in endometriosis patients ^77^, and peripheral blood mononuclear cells of multiple sclerosis patients ^78^. Conversely, it is downregulated in the peripheral blood mononuclear cells of Sjögren’s syndrome patients ^79^. These results suggest that this gene being switched on versus off may predispose individuals to certain diseases while protecting them against others. This balance between susceptibility and protection could explain why both high-expression and low-expression states are maintained in the population at comparable frequencies.

Thirdly, the bimodality of *NPIPA5* (**Figure S2G**), too, can be explained by a single eQTL. The T allele of the SNV rs3198697 is associated with *NPIPA5* being switched on across tissues, while the C allele is associated with the gene being switched off. *NPIPA5* has been reported as one of the top differentially expressed genes among patients with multiple sclerosis in both blood and brain ^80^. Moreover, this study ^80^ showed that this gene is co-expressed in blood and brain. Here, we have shown that this gene is switch-like and that the co-expression of *NPIPA5* is not restricted to blood and brain but extends to all pairs of tissues.

Lastly, a single eQTL can explain the bimodality of a member of the PKD1 gene family in cluster 2A, *PKD1P5* (**Figure S2I**). For *PKD1P5*, the C allele of the SNV rs201525245 is associated with the gene being switched on, while the G allele is associated with the gene being switched off.

**Figure S2.**
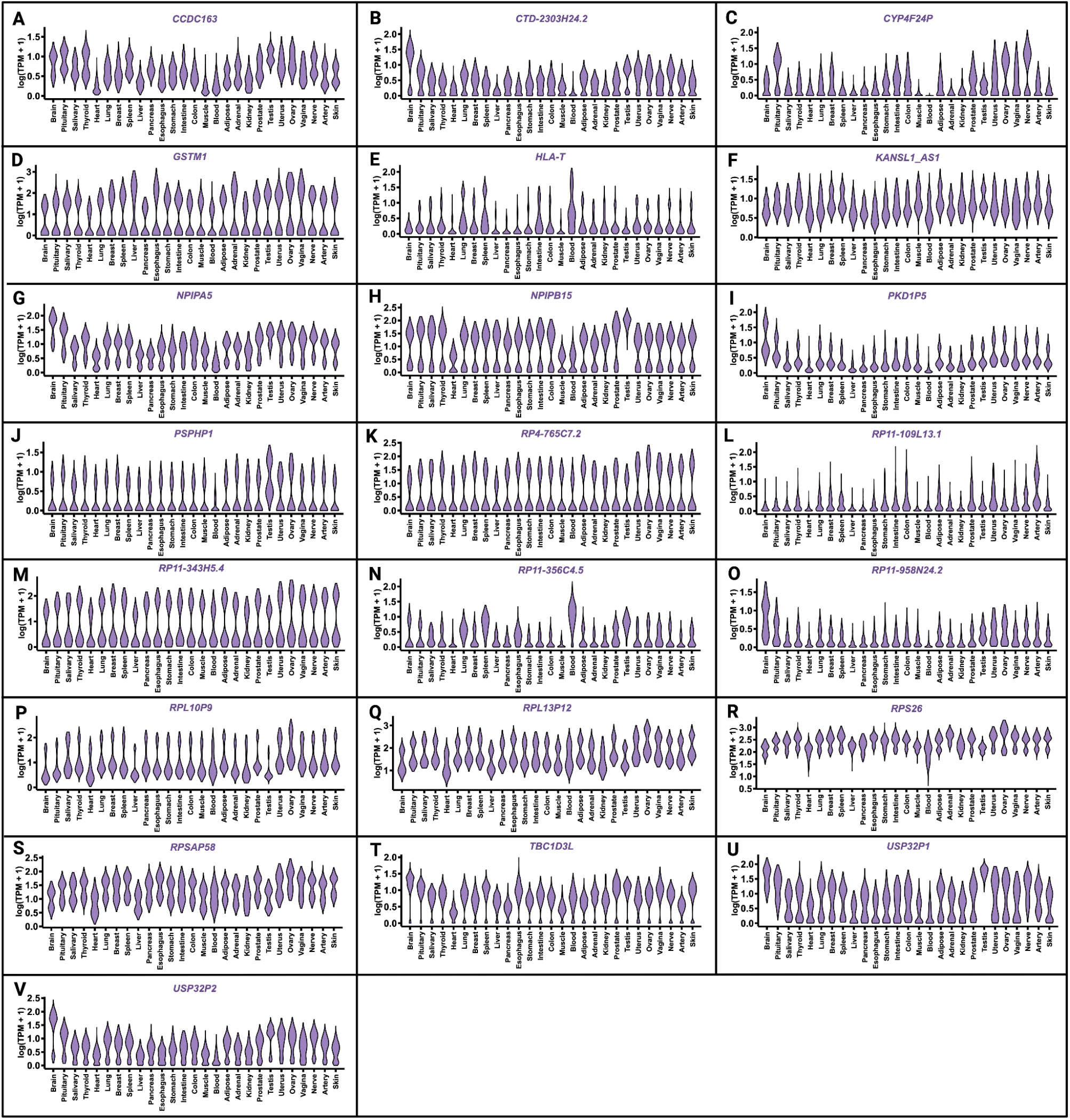
Violin plots for expression level distributions of switch-like genes in cluster 2A.

**Figure S3.**
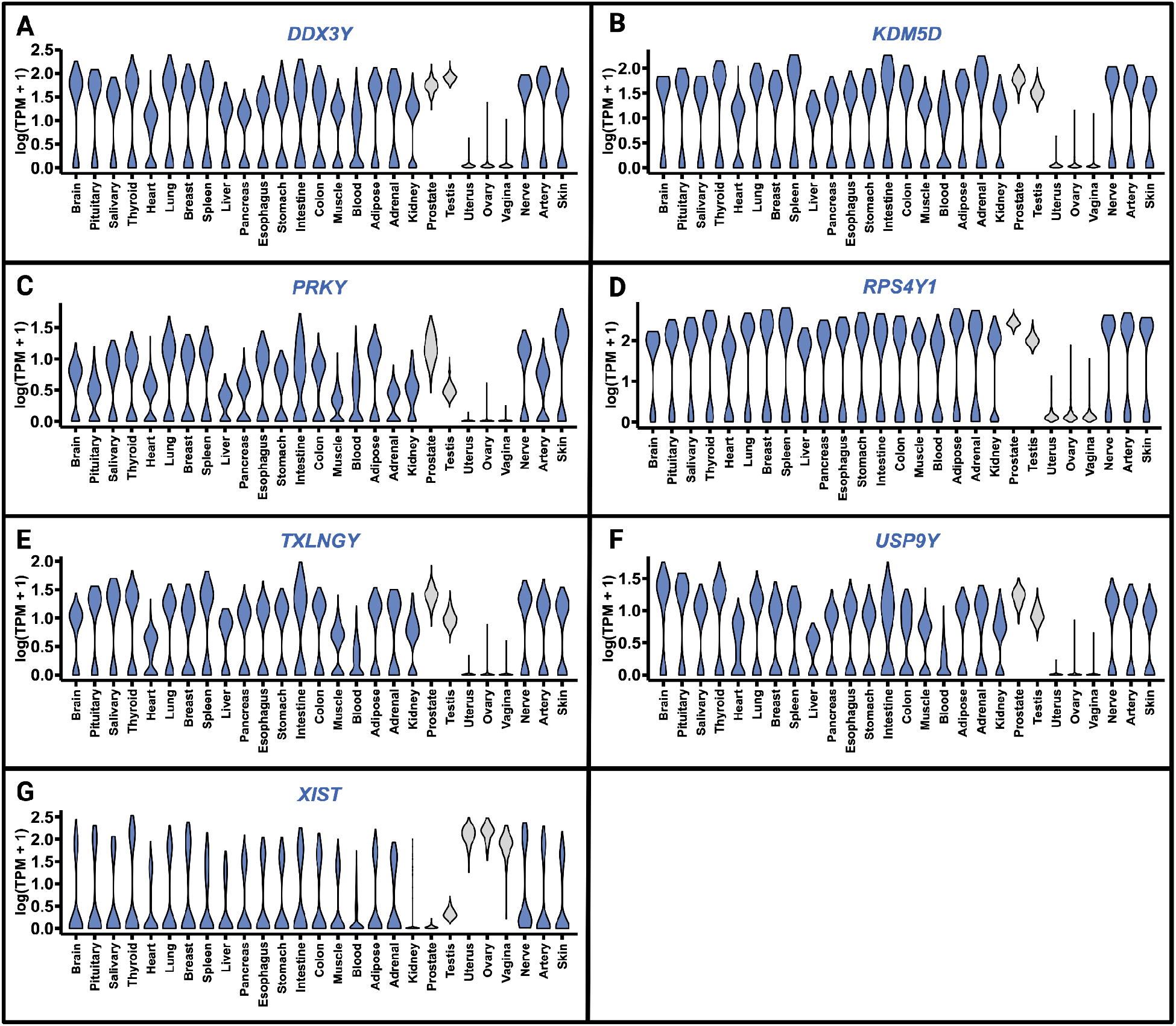
Violin plots for expression level distributions of switch-like genes in cluster 2A.

### Conceptual issues regarding bimodal expression distributions driven by genetic polymorphisms

In the main text, we claimed that genetic polymorphisms drive the bimodal expression of universally switch-like genes in cluster 2A. For a polymorphism with two alleles (*A* and *a*), there are three possible genotypes (*aa, Aa, and AA*). Since each of the three genotypes can lead to three different expression levels, we expect expression distributions of a cluster-2A gene to have three modes. This leads to the question: Why do we not see trimodal, as opposed to bimodal, expression distributions for genes in cluster 2A? To answer this question, we develop the following frameworks. Let us assume that a genetic polymorphism exists with two alleles, *A* and *a*, with frequencies *p*_*A*_ and (1−*p*_*A*_), respectively. The three genotypes for this polymorphism, *aa, Aa*, and *AA*, lead to three different expression states (TPM levels) for the gene with averages *μ*_*aa*_, *μ*_*Aa*_, and *μ*_*AA*_, respectively. Let us also assume that the Hardy-Weinberg equilibrium holds for this locus. Then, the frequency of *aa* = (1−*p*_*A*_)^2^, the frequency of *Aa =* 2*p*_*A*_ (1−*p*_*A*_), and the frequency of *AA* = *p*_*A*_^2^. We assume that *μ*_*aa*_ & *μ*_*Aa*_ & *μ*_*AA*_. Next, we define a dominance coefficient 0 ≤ *α* ≤ 1 by,

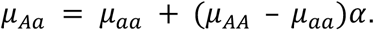

If we define the ratio *R* by

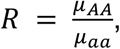

then, we obtain

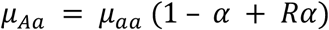

and

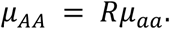

We can then divide individuals into three groups depending on their genotypes. Let us assume that the coefficient of variation (CV) of expression is the same for each genotypic group. Then, we can model the TPM value of this gene in a given individual a normal random variable with:

1. mean = *µ*_*aa*_ and standard deviation = CV x *μ*_*aa*_ if the genotype is *aa*;
2. mean = *µ*_*Aa*_ and standard deviation = CV x *μ*_*Aa*_ if the genotype is *Aa*; and
3. mean = *µ*_*AA*_ and standard deviation = CV x *μ*_*AA*_ if the genotype is *AA*.

The value of *µ*_*aa*_ is irrelevant for gauging the effect of polymorphisms on the shape of the expression level distributions. Therefore, we set *µ*_*aa*_ = 1.

Under these mathematical assumptions, we performed simulations using 36 distinct models. These models vary by four parameters: *p*_*A*_ ∈ {0.05, 0.1, 0.5}, CV ∈ {0.1, 0.3}, *R* ∈ {10, 1000}, and *α* ∈ {0.2, 0.5, 0.8}. For each model, defined by a unique combination of the values of these four parameters, we performed a two-step sampling procedure. First, we obtained a random sample of 500 genotypes, based on *p*_*A*_ and the Hardy-Weinberg equilibrium. Next, for each of the 500 genotypes sampled, we sample a TPM value from the normal distribution corresponding to that genotype. Thus, for each of the 36 models, we simulated 500 TPM values. We present these values as histograms with and without log transformation. The results for *p*_*A*_ = 0.05, *p*_*A*_ = 0.1, and *p*_*A*_ = 0.5 are shown in **Figure S4, Figure S5**, and **Figure S6**, respectively. These simulations help us answer our question we first asked: Why do we not see a trimodal distribution if a genetic polymorphism drives expression-level variability in a gene?

Firstly, even when the minor allele (*A*) frequency is not low (e.g., 10%), the frequency of the genotype *AA* is still quite low (e.g., 1%). Therefore, the third peak is not always conspicuously visible. We see this in all models with *p*_*A*_ = 0.05 and *p*_*A*_ = 0.1 (**Figures S4** and **S5**), regardless of CV, *R*, and *α* values. At higher allele frequencies (e.g., 50%), the effect of the remaining parameters becomes more apparent. **Figure S6** shows that a higher dominance coefficient *α* makes the expression level distribution more bimodal. By contrast, a lower dominance coefficient *α* makes the expression level distribution more trimodal. The lack of observed trimodality in the GTEx data may suggest that expression levels of switch-like genes tend to be more dominant than additive with regard to causal genetic polymorphisms. Secondly, greater variation (CV) in the data can also obscure the third peak. For example, by comparing **Figure S6B** to **Figure S6H**, we find that increasing the CV can change the distribution from being trimodal to bimodal when the other parameters are held constant. However, *R* does not seem to have much effect on whether the expression level distribution is bimodal or trimodal.

**Figure S4.**
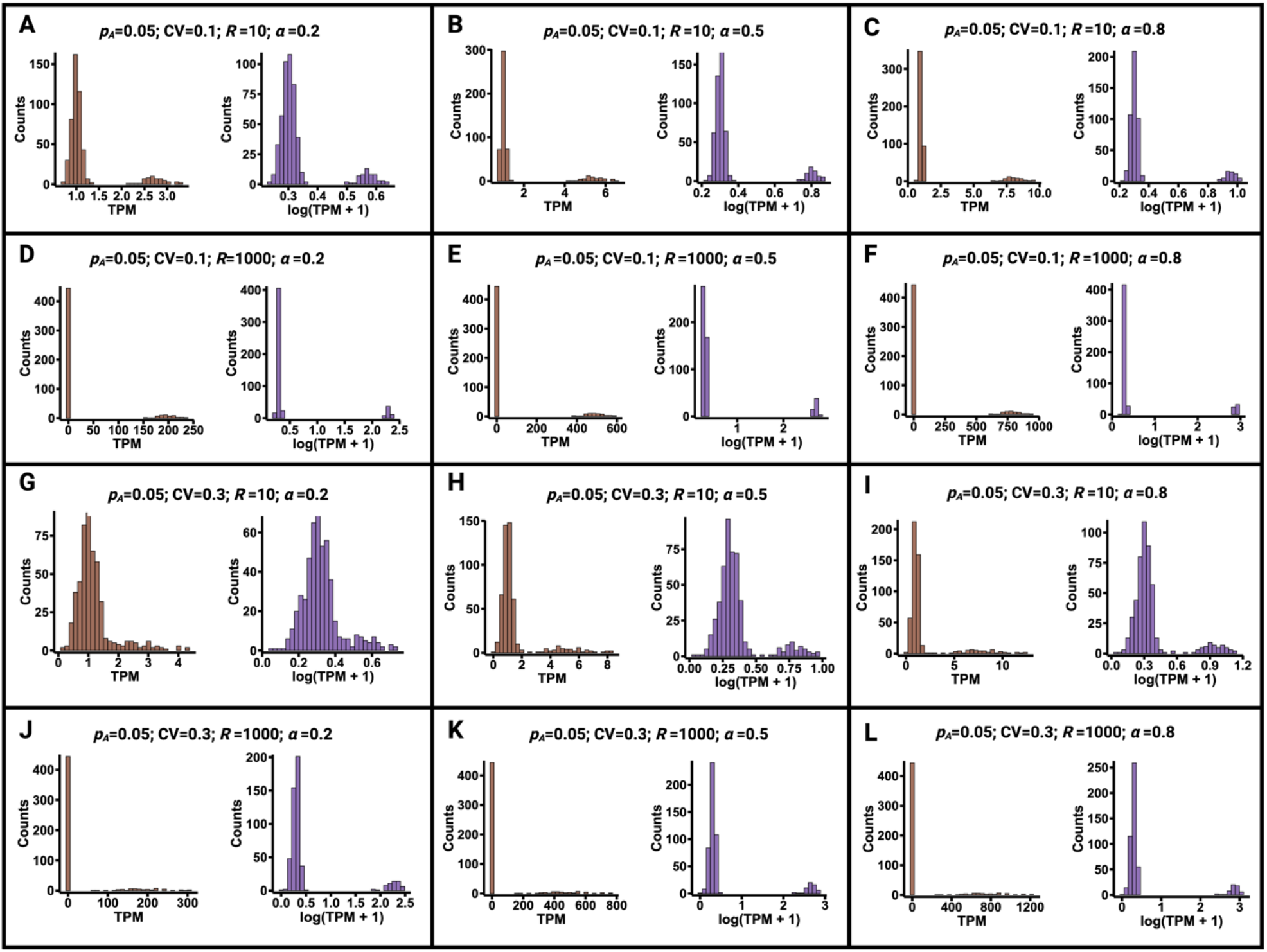
TPM simulations for a hypothetical gene whose expression is driven by a genetic polymorphism with an allele frequency of 5%.

**Figure S5.**
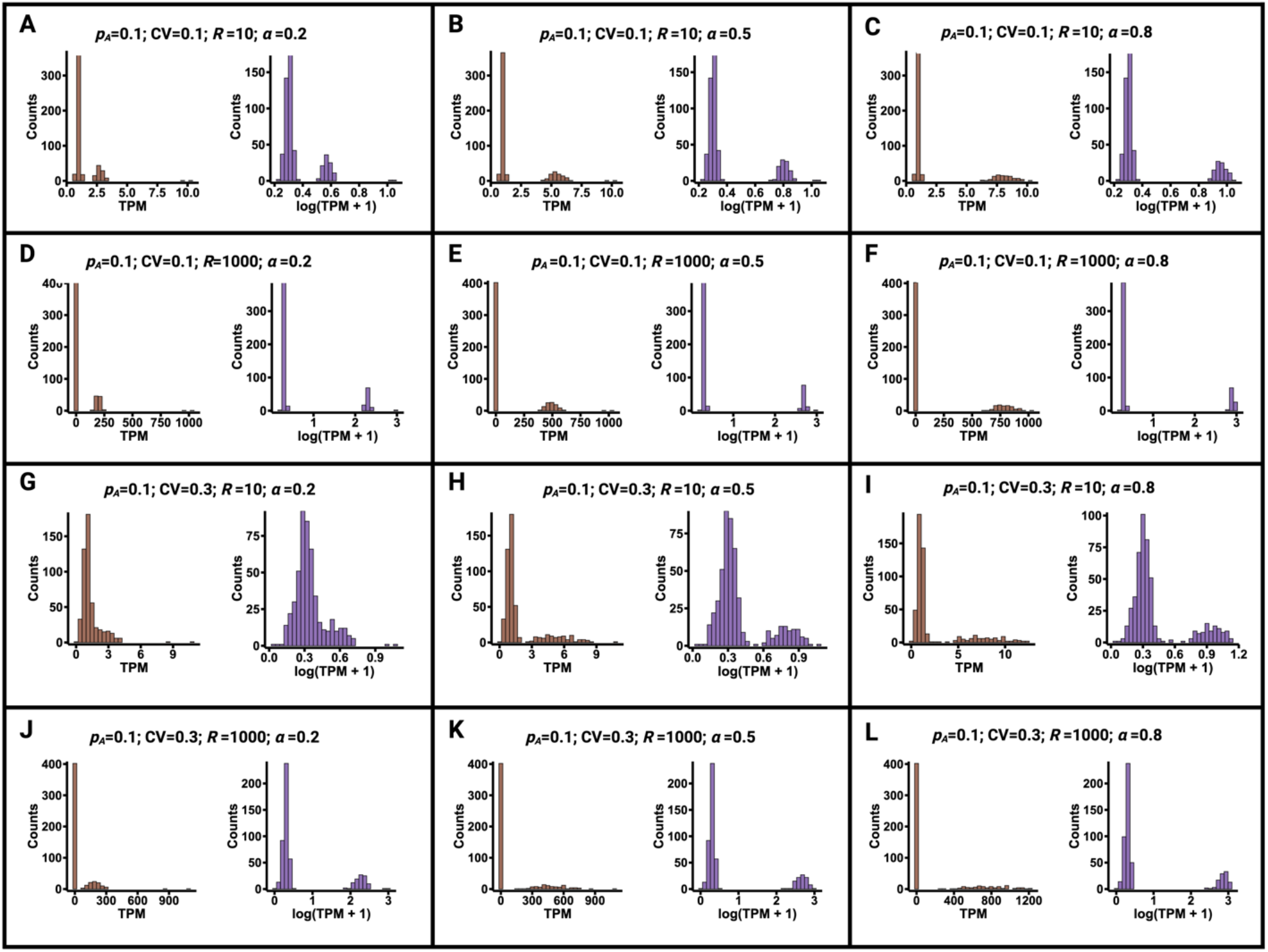
TPM simulations for a hypothetical gene whose expression is driven by a genetic polymorphism with an allele frequency of 10%.

**Figure S6.**
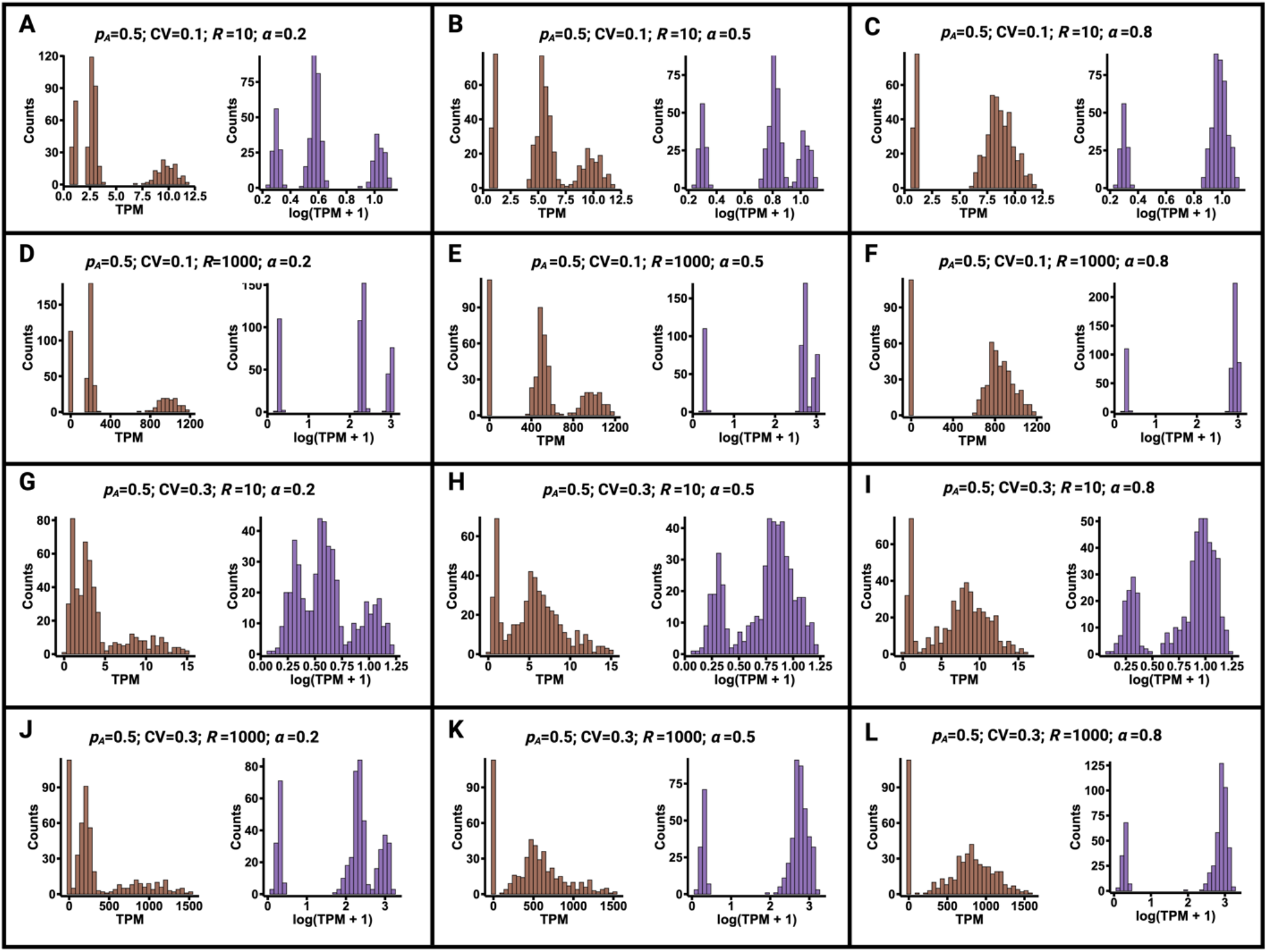
TPM simulations for a hypothetical gene whose expression is driven by a genetic polymorphism with an allele frequency of 50%.

### Tissue-specific switch-like genes

We divided switch-like genes into three clusters in the space spanned by the first two principal components (**Figure 2A**). While we said that genes in cluster 1 **(Figure 2A-B)** are tissue-specific switch-like genes, manual inspection reveals this is not true for all genes in cluster 1. In particular, the transcript ENSG00000273906 coming from chr Y was labeled cluster 1 by hierarchical clustering even though it is universally switch-like in tissues common to both sexes. Indeed, we removed all chr-Y genes from our analyses of genuine cluster-1 genes. Other cluster-1 genes bimodally expressed in a large number of tissues lie on the autosomes. For example, *CLPS, PRSS1, CELA3A*, and *CELA3B*, despite having low overall tissue-to-tissue co-expression, are bimodally expressed across tissues. Indeed, we have shown previously that *CELA3A* and *CELA3B* have a shared regulatory architecture in the pancreas ^81^.

### Controlling for confounders

We removed cluster-1 genes affected by confounders in each tissue using an approach outlined in **Methods**. Here, we present the number of genuine cluster-1 genes versus those affected by confounders in **Figure S7**. In particular, we show that the cluster-1 genes in the colon and the intestine are particularly prone to being affected by confounding factors. We also present in **Figure S8** examples of genes whose bimodal expression in specific tissues is correlated with variation in the sample ischemic time distribution.

**Figure S7.**
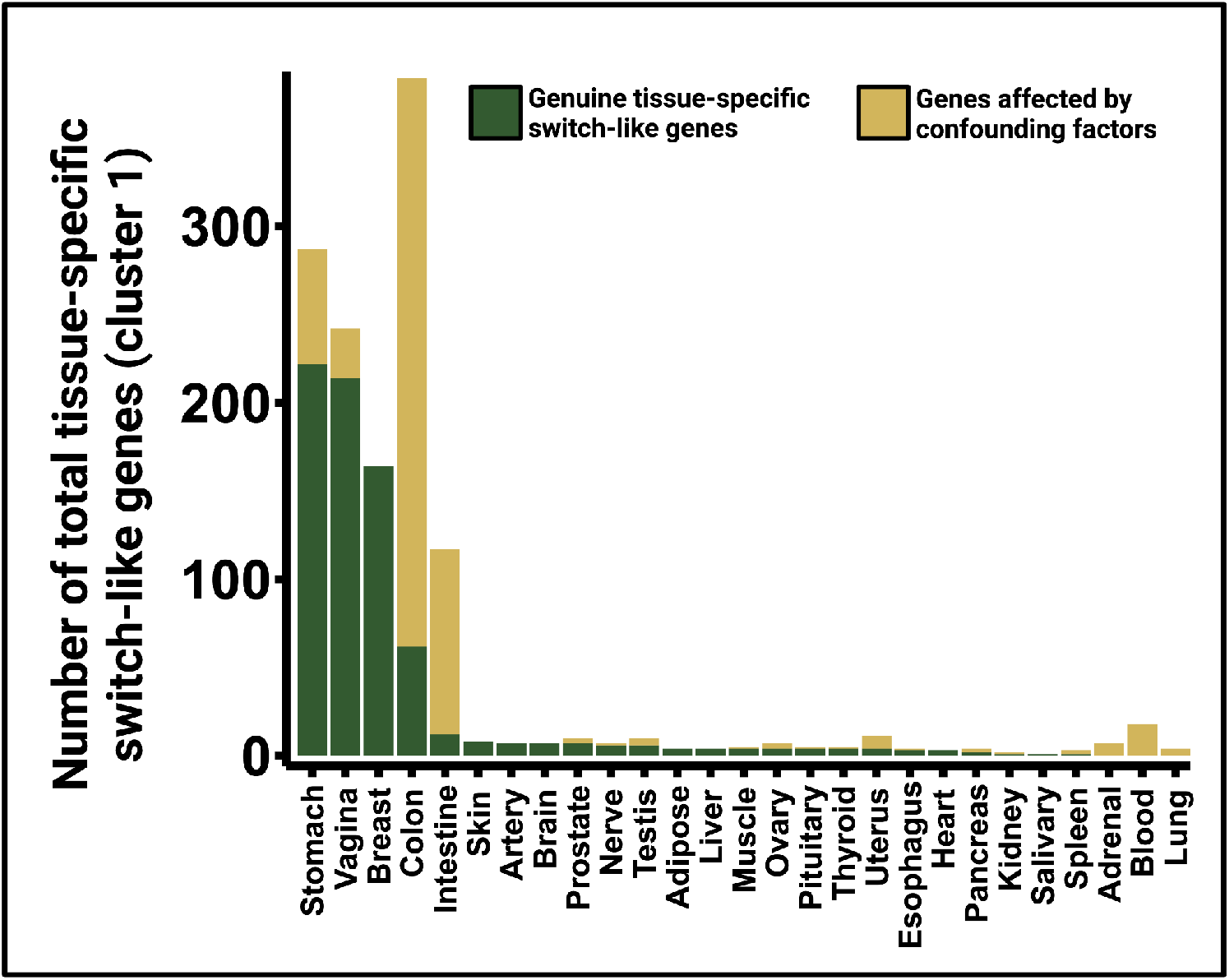
Switch-like genes in cluster 1 that are genuine versus those affected by confounders.

**Figure S8.**
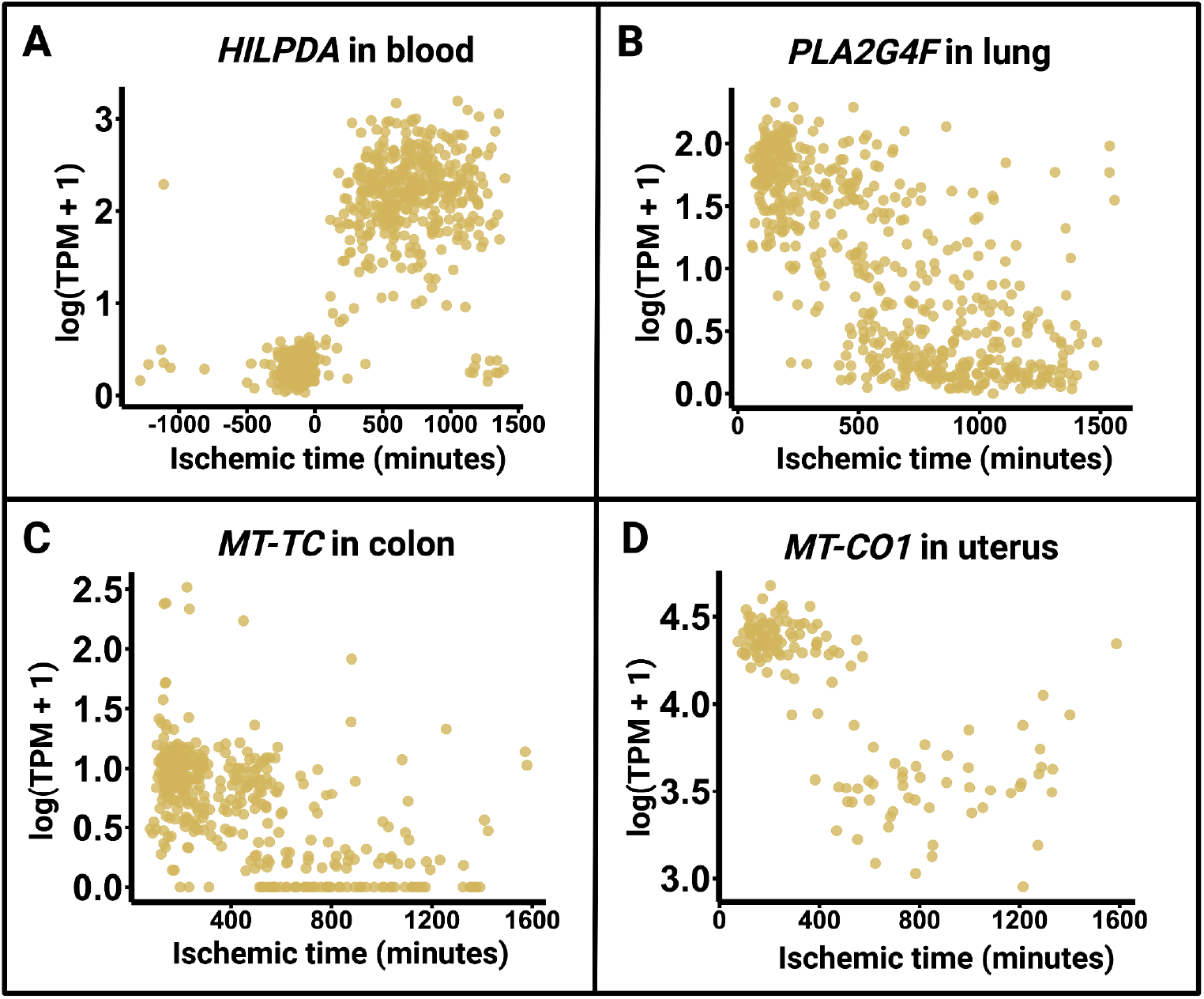
Examples of cluster-1 genes affected by confounders. Their bimodal distribution is caused by ischemic time (a confounding factor).

### The copy number variation at the *PGA3* locus does not affect the gene’s expression levels

*PGA3* exhibits a high copy number variation among humans ^82^, but the copy number seems to have no impact on *PGA3* expression, at least in cancer samples ^83^. The bimodal expression of PGA3 in the stomach is likely not due to its copy number variation. This is because PGA3’s expression in the stomach is highly correlated with other tissue-specific genes in the stomach. The only way in which a copy number-driven bimodality of PGA3 could be correlated with other switch-like genes is if the product of PGA3 was regulating the correlated genes. Without this evidence, we surmise that the copy number variation at the PGA3 locus does not affect the gene’s expression levels, at least in the stomach.

**Table S1. A list of tissues used in this study along with the number of individuals for each tissue.**

**Table S2. A list of 1**,**013 switch-like genes**.

**Table S3. Tissue-to-tissue co-expression (Pearson’s correlation) for all genes across 310 tissue-tissue pairs**.

**Table S4. Results from principal component analysis on tissue-to-tissue co-expression data for all genes**.

**Table S5. Results from principal component analysis on tissue-to-tissue co-expression data for only switch-like genes**.

**Table S6. Correlation between gene expression levels and confounding factors for switch-like genes**.

**Table S7. Gene-to-gene co-expression of genuine tissue-specific switch-like genes in the stomach, vagina, breast, and colon**.

**Table S8. Analysis of sex bias among genuine tissue-specific switch-like genes**.

